# Lifelong regeneration of cerebellar Purkinje neurons after induced cell ablation in zebrafish

**DOI:** 10.1101/2022.05.10.491347

**Authors:** Sol Pose-Méndez, Paul Schramm, Barbara Winter, Jochen C. Meier, Konstantinos Ampatzis, Reinhard W. Köster

## Abstract

Zebrafish have an impressive capacity to regenerate neurons in the central nervous system. However, regeneration of the principal neuron of the evolutionary conserved cerebellum, the Purkinje cell (PC), is believed to be limited to developmental stages based on invasive lesions. In contrast, non-invasive cell type specific ablation by induced apoptosis closer represents processes of neurodegeneration. We demonstrate that the ablated larval PC population entirely recovers in number, quickly reestablishes electrophysiological properties and properly integrates into circuits to regulate cerebellum-controlled behavior. PC progenitors are present in larvae and adults, and PC ablation in adult cerebelli results in an impressive PC regeneration of different PC subtypes able to restore behavioral impairments. Interestingly, caudal PCs are more resistant to ablation and regenerate more efficiently, suggesting a rostro-caudal pattern of de- and regeneration properties. These findings demonstrate that the zebrafish cerebellum is able to regenerate functional PCs during all stages of the animal’s life.

## Introduction

Regeneration of neurons in the central nervous system of mammals has been demonstrated to occur, but it is rare and limited to few brain areas (Iismaa et al., 2018). In contrast, in some amphibians and fish, regeneration in almost any part of the central nervous system is evident and occurs not only during development or in juveniles, but also during adulthood (Kizil et al., 2012; Lust and Tanaka, 2019; Tanaka and Reddien, 2011; Zambusi and Ninkovic, 2020). Hence, evolutionary conserved fish brain compartments represent a valuable testing ground for characterizing neuronal regeneration in vertebrates.

The zebrafish and mammalian cerebellum share nearly all neuronal cell types with their circuitry, physiology and function being conserved (Hashimoto and Hibi, 2012; Hibi et al., 2017; Koyama et al., 2021; Matsuda et al., 2017; Volkmann et al., 2010). Furthermore, clarification of the developmental origin and differentiation program of zebrafish cerebellar neurons, their physiology and functional contribution to locomotor control, motor learning and socio-emotional behavior has been revealed, and these studies have laid a solid foundation for regeneration surveys of cerebellar neurons (Chang et al., 2021b, 2020; Harmon et al., 2017; Kidwell et al., 2018; Knogler et al., 2019, 2017; Koyama et al., 2021; Markov et al., 2021; Matsuda et al., 2017; Matsui et al., 2014; Rieger et al., 2009; Volkmann et al., 2008).

Based on the extensive adult neurogenesis and regenerative capacity, it is widely believed that all neurons within the zebrafish central nervous system are able to regenerate (Kroehne et al., 2011). Yet, in elegant studies this long-held classical view has recently been upset for highly conserved key cerebellar neurons, the Purkinje cells (PCs) (Kaslin et al., 2017).

PCs constitute the principal information processing and output neurons of the cerebellum. Hence, PC degeneration in humans results in severe neurological symptoms like ataxia, altered locomotor activity, but also anxiety and social disabilities. These diseases can be genetically modeled in zebrafish PCs (Elsaey et al., 2021; Hsieh et al., 2020; K. Namikawa et al., 2019; Kazuhiko Namikawa et al., 2019; Watchon et al., 2017), but it has remained elusive so far whether such PC degeneration can be counteracted by regeneration to slow down and mitigate disease progression.

During embryonic stages the entire cerebellar primordium including PCs regenerates after surgical removal, due to repatterning processes in the hindbrain (Köster and Fraser, 2006). Also, at larval and juvenile stages, the cerebellum recovers from local wounding and regenerates PCs (Kaslin et al., 2017). But this PC regeneration capacity declines and is lost at 3 months of age, which has been explained by the exhaustion of proper PC progenitors, while progenitors of other cerebellar neuronal cell types continue to undergo neurogenic divisions (Hentig et al., 2021; Kaslin et al., 2017). These findings comprehensively suggest that the regeneration capacity of zebrafish PCs is ambiguous, regenerating efficiently during larval and juvenile stages as expected for teleostean neurons, but lacking such a capacity during adulthood resembling mammalian PCs (Kaslin et al., 2017).

Yet, larval PC regeneration upon local wounding has been based on the appearance of new PCs, but this could represent ongoing growth of the PC population rather than a regenerative response. In addition, PCs considered to have regenerated cannot be distinguished from non-injured neighbors, therefore the proper physiological integration of regenerated larval PCs into existing circuitry could not be revealed, therefore also in larvae functional PC regeneration is still at question.

Conversely, injuring the adult cerebellum may mask the ability to regenerate PCs. Extensive surgical ablations remove important progenitors and the cellular and extracellular environment likely required for PC regeneration. Local traumatic injuries instead may be counteracted by plasticity of neighboring PCs. In addition, acute traumatic cerebellum injuries affect numerous cell types beyond PCs in a defined area, and if invasive, induce wound healing that could influence regeneration (Hentig et al., 2021; Kaslin et al., 2017).

Instead, degenerative diseases of PCs affect the entire population of a single cell type by non-invasive cell death mechanisms. Likely, these principally different situations of PC loss pose different demands and constraints on regeneration. Therefore, the ability of larval and adult PCs to regenerate is still enigmatic.

We have recently introduced transgenic PC-ATTAC^TM^ zebrafish, in which PCs can be selectively ablated by apoptosis through non-invasive administration of Tamoxifen (Weber et al., 2016). This model combines several advantages to address the open question of the occurrence of functional PC regeneration: **a)** Ablating nearly the entire PC population uncouples plasticity from regeneration and creates a high regenerative demand; **b)** PC death by apoptosis and microglia phagocytosis (Weber et al., 2016) prevents leakage of PC contents into the extracellular environment and resembles disease-associated neurodegeneration and; **c)** Fluorescent protein expression in PCs allows not only for quantifying ablation and recovery efficiencies, but also for investigating physiological integrity of regenerating PCs (Hsieh et al., 2014) and relating behavioral recovery to the extent of PC population regrowth (Champagne et al., 2010; Ying-Yu Huang, 2008); **d)** Age-independent Tamoxifen treatment allows to study PC regeneration in larvae, juveniles and adults under equivalent conditions. Using this PC specific ablation approach, we have readdressed the open question of PC regeneration by combining *in vivo* imaging, electrophysiology and behavioral analysis in both the larval and the adult zebrafish cerebellum.

## Results

### Recovery of larval Purkinje cells (PCs) after induced non-invasive ablation

Carriers of the stable transgenic zebrafish strain PC-ATTAC^TM^ express a membrane targeted red fluorescent protein (FyntagRFP-T) as reporter together with a Tamoxifen-inducible Caspase 8 selectively in postmitotic cerebellar Purkinje neurons (Figure 1A) (Weber et al., 2016). Between 4-6 days post-fertilization (dpf) the PC layer in zebrafish is established and matures (Hamling et al., 2015; Kazuhiko Namikawa et al., 2019). Incubation of PC-ATTAC^TM^ larvae between 4 and 6dpf with 4-hydroxy-tamoxifen (4-OHT) overnight induced apoptosis in the majority of PCs (85-95%) leaving red fluorescent apoptotic debris behind, which progressively disappeared by microglial phagocytosis within 3-5 days post-treatment (dpt) (Weber et al., 2016). Since ethanol (EtOH) was used as solvent for 4-OHT stock solutions, EtOH-treated (0.4% final concentration) specimens were used as controls, which did not show any sign of PC death (Figure 1B).

**Figure 1:**
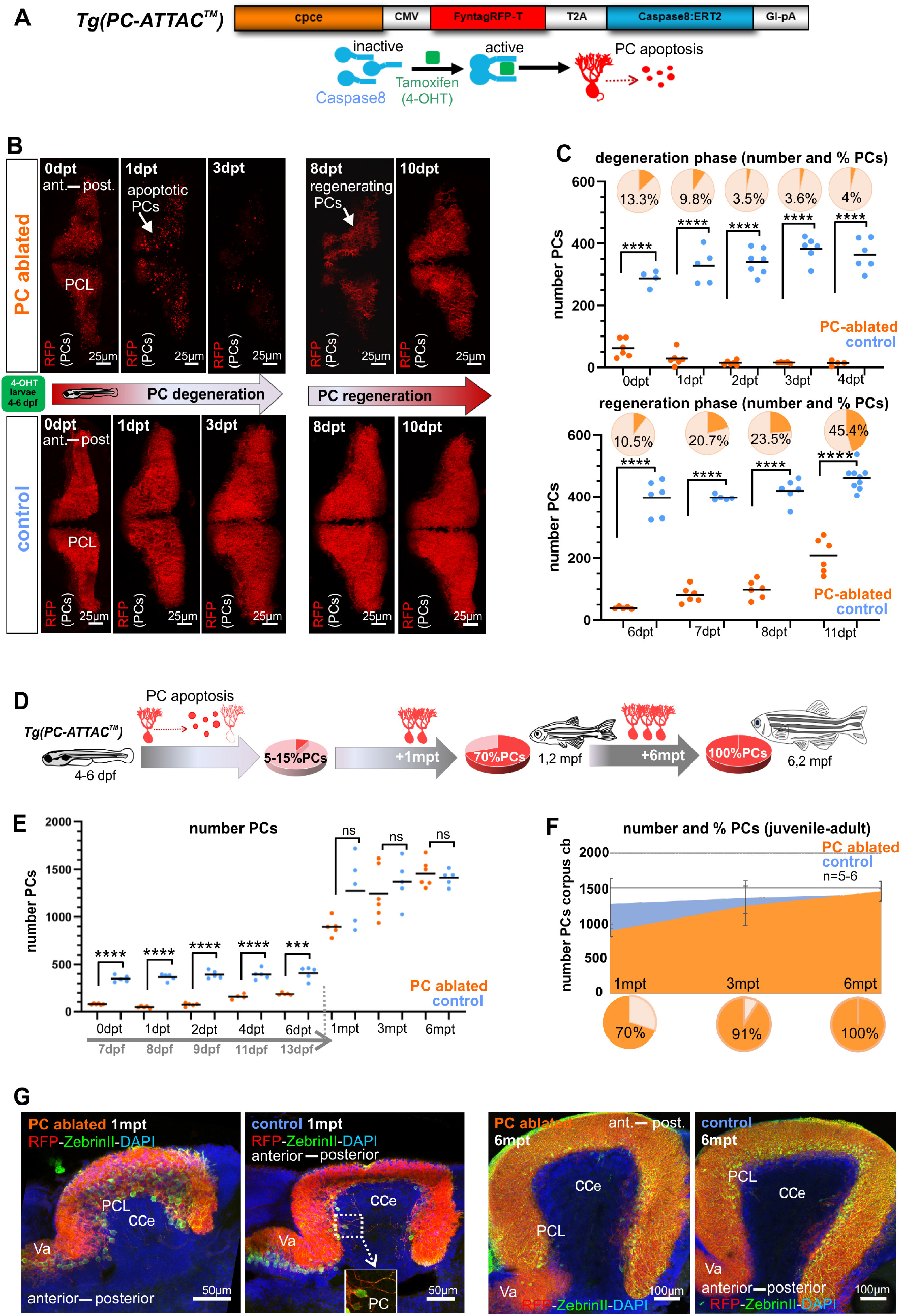
Induced PC ablation in zebrafish larvae: time course of degeneration and regeneration. **(A)** Construct used to generate the Tg(PC-ATTAC^TM^) transgenic line, modified from (Weber et al., 2016). **(B)** Images of the PC layer (fyn-tagRFP-T fluorescence expressed in mature PCs) after induced PC ablation in larvae at 4-6 dpf monitoring 10 days after 4-OHT treatment. **(C)** Number and percentage of PCs in ablated (4-OHT) vs. control (EtOH) larvae. **(D-G)** Quantitative monitoring of PC regeneration after induced PC ablation in larvae. D-F) Numbers and percentage of PCs until 6 mpt. G) Images of PCs on sagittal cerebellar sections after immunostaining with anti-tagRFP and anti-ZebrinII antibodies, comparing ablated and control groups at 1 and 6 mpt. **Figure Supplement 1A-D.** Quantification of cell proliferation by BrdU-labeling after PC ablation. **Source data 1.** PC quantification for Figure 1C, D, F.

About 5-7dpt first red fluorescent PCs reappeared, and already after about 10dpt half of the PC population compared to controls was replenished (Figure 1B, C). This demonstrates that during larval stages the PC population can quickly recover from a nearly complete removal.

### Ablated larval PC population regenerates completely

The partial PC regeneration observed within a few days could either represent a regenerative response, the continuation of PC layer development or a combination of both. Only in the latter case the PC population would regain its full size compared to controls. Therefore, PC numbers were quantified from 1 to 6 months post-treatment (mpt) (Figure 1D-G). Compared to PCs found in non-ablated controls, numbers of PCs in specimens after initial PC ablation reached 70% at 1mpt, 91% at 3mpt, and 100% at 6mpt, respectively (Figure 1D-F). Since PCs ablated during larval development (ca. 400) amount to about one quarter of the adult PC population at 6 months (ca 1.500), an obvious reduction in PC population size would be expected, if the ablated PCs were not replaced, but such a reduction in PC numbers was not observed. Instead, a full recovery of PCs was evident at 6mpt.

In addition, no apparent anatomical differences of the cerebellar cortex at 1mpt and 6mpt could be observed by immunohistochemistry against the PC-specific antigen ZebrinII on sagittal sections from PC-ablated and control specimens (Figure 1G). These findings corroborate the complete restoration of the ablated larval PC population (Figure 1D-G).

To obtain further insight into the temporal dynamics of PC replacement, proliferating cells were labeled by Bromodeoxyuridine (BrdU) treatment for 24 hours at different time points post-ablation followed by their immunohistochemical detection and quantification. During the first seven days post-ablation no significant increase in proliferating cell numbers could be observed in PC ablated larvae compared to vehicle treated controls (**Figure 1 - figure supplement 1A-D**). This showed that PC regeneration was not initiated by a burst in cell proliferation, but occurred continuously by slowly adding surplus PCs over time until the full PC population size was reestablished 6 months later.

### Progenitor cells of regenerating PCs

Since radial glia are considered one of the potential sources for neuronal regeneration, we crossed carriers of the transgenic Tg(*gfap*:EGFP) strain with green fluorescent radial glia cells into the PC-ATTAC^TM^ background. Yet, after PC ablation none of the reappearing PCs expressed both fluorescent reporters (Figure 2A), nor did the radial glia cell population increase significantly (Figure 2B). This makes a contribution of radial glia to larval PC regeneration unlikely, although it cannot be excluded that *gfap*-enhancer driven EGFP expression could have been lost by the time regenerating PCs differentiate.

**Figure 2:**
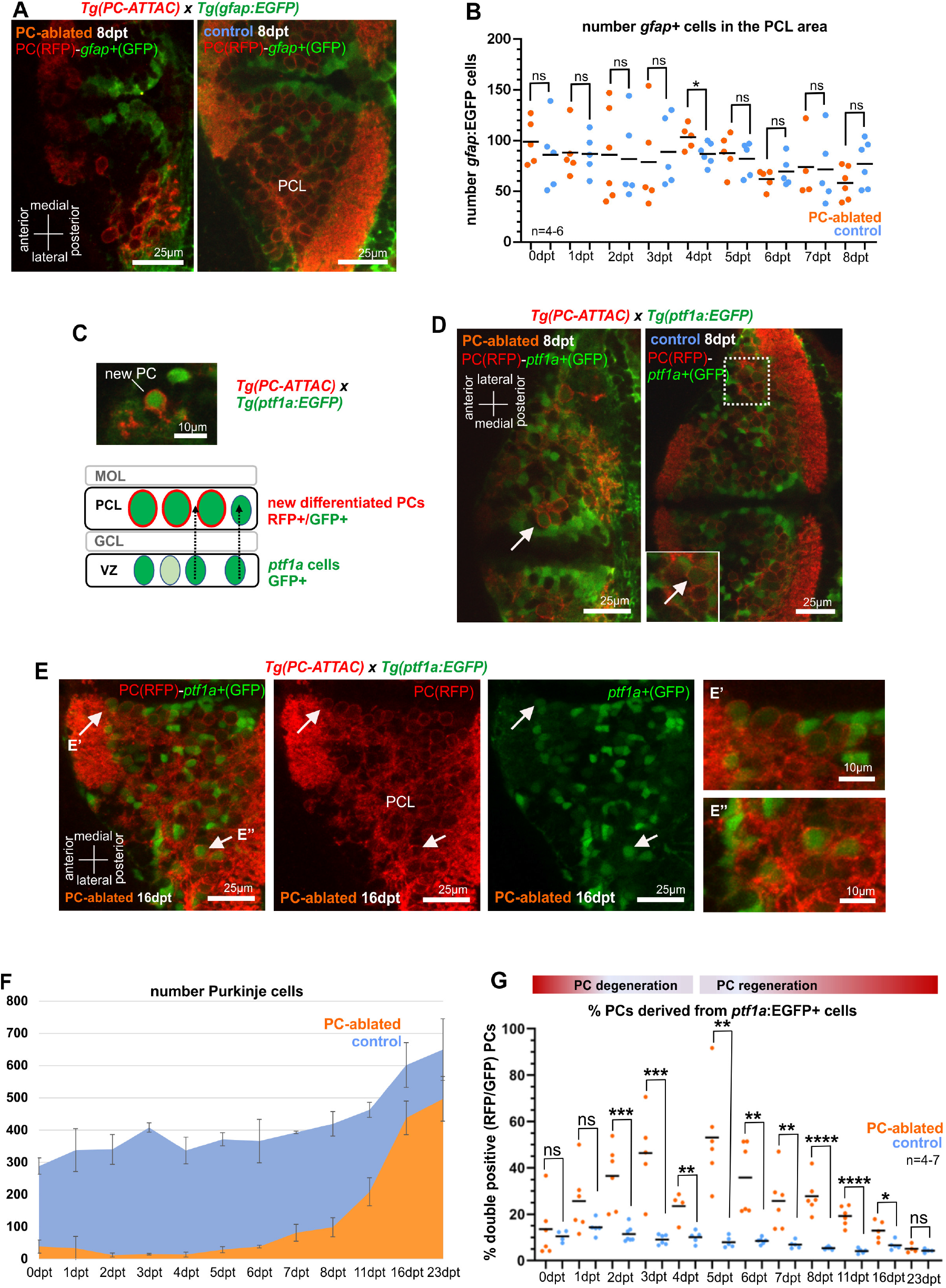
Cellular analysis of potential progenitors of regenerating PCs. **(A)** Images of larval cerebellum of the double transgenic line Tg(PC-ATTAC)/Tg(*gfap*:EGFP) after induced PC ablation. **(B)** Number of gfap^+^ cells throughout the PC layer area during PC degeneration and beginning of regeneration. **(C)** Illustration of new PC development from *ptf1a*^+^-progenitors. **(D, E)** Images of larval cerebellum of the double transgenic line Tg(PC-ATTAC)/Tg(*ptf1a*:EGFP) after induced PC ablation, revealing double positive cells (arrows). **(F, G)** Average of PC numbers (F), and percentage of PCs showing GFP fluorescence (double positive cells, G) during degeneration and regeneration of PCs. The red fluorescent protein from the PC-ATTAC strain is exclusively expressed in the cell membrane, while EGFP from the *ptf1a-* and *gfap*-reporter strains localizes to the cytoplasm. GFP and RFP were enhanced by fluorescence immunohistochemistry in 2A, C-E. **Figure Supplement 1E-F.** *ptf1a*:GFP positive PCs and absolute number of *ptf1a*:GFP cells in the cerebellum after PC ablations compared to controls. **Figure Supplement 1G.** Cumulative BrdU labeling of regenerating PCs. **Source data 1.** Quantification of *gfap*-expressing cells for Figure 2B and quantification of *ptf1a*:GFP-expressing PC-ATTAC cells for Figure 2G.

As an alternative, ventricular zone (VZ) derived neuroepithelial cells were tested that give rise to GABAergic cerebellar neurons. These cells likely originate from neural stem/progenitor cells (NSPCs) (Chang et al., 2021a; Jung et al., 2013) and express the green fluorescent protein in the transgenic strain Tg(*ptf1a*:EGFP) (Bae et al., 2009). Indeed, double fluorescent PCs in Tg(*ptf1a*:EGFP) larvae in the PC-ATTAC^TM^ background were observed following PC ablation (Figure 2C-E) and were significantly increased in their ratio compared to only red fluorescent PCs for about two weeks starting two days after ablation (Figure 2F, G).

The ratio of double fluorescent PCs is in favor of newly differentiated PCs, as EGFP expression by the *ptf1a*-enhancer is only transient. Mature PCs in control specimens therefore exceed mature PC numbers in larvae with active PC regeneration (**Figure 2 - figure supplement 1E**). The absolute number of *ptf1a*:EGFP expressing cells in the cerebellum was not significantly higher in PC-ablated PC-ATTAC^TM^ larvae compared to controls (**Figure 2 - figure supplement 1F**) consistent with the BrdU labeling results. In addition, a small increase in green fluorescent PC progenitor cells may be further masked as *ptf1a*:EGFP expressing cells contribute to several neuronal lineages of the cerebellum (Bae et al., 2009). In conclusion, larval PC regeneration is driven by a small increase of VZ-derived *ptf1a*-expressing neuroepithelial cells, the natural PC progenitor population, and these cells may not express the radial glia marker *gfap* or have lost *gfap*-enhancer-driven EGFP-expression at the time of PC differentiation.

Interestingly, both double fluorescent recently differentiated PCs (Figure 2D, E) and regenerating PCs marked by cumulative BrdU labeling (**Figure 2 - figure supplement 1G**) were not randomly distributed, but clustered loosely in two domains along the dorsal midline and in lateral parts of the cerebellum, respectively. This suggests that the VZ is spatially patterned like the adjacent germinal zone of the upper rhombic lip (Volkmann et al., 2008).

### Physiology of regenerating PC circuitry

To obtain a reference for circuit reestablishment by regenerating PCs, we first analyzed the electrophysiological properties of PCs in 4-21dpf old larvae of the PC reporter line Tg(−7.5ca8:GFP)^bz12^ under resting conditions by patch-clamp recordings in attached cell configuration (Figure 3A-E). These studies confirmed that differentiating PCs require about two days to establish a stable firing pattern of a mature PC with an average firing frequency between 8-10Hz, displaying high-frequency bursts of over 30Hz and a ratio for complex spikes to simple spikes cs/ss of 0.04 (Harmon et al., 2017; Hsieh et al., 2014; Scalise et al., 2016; Sengupta and Thirumalai, 2015). First PCs born from 4-6dpf forming the initial PC layer reach a plateau of electrophysiological maturity around 8dpf (**Figure 3 - figure supplement 2A-C**).

**Figure 3:**
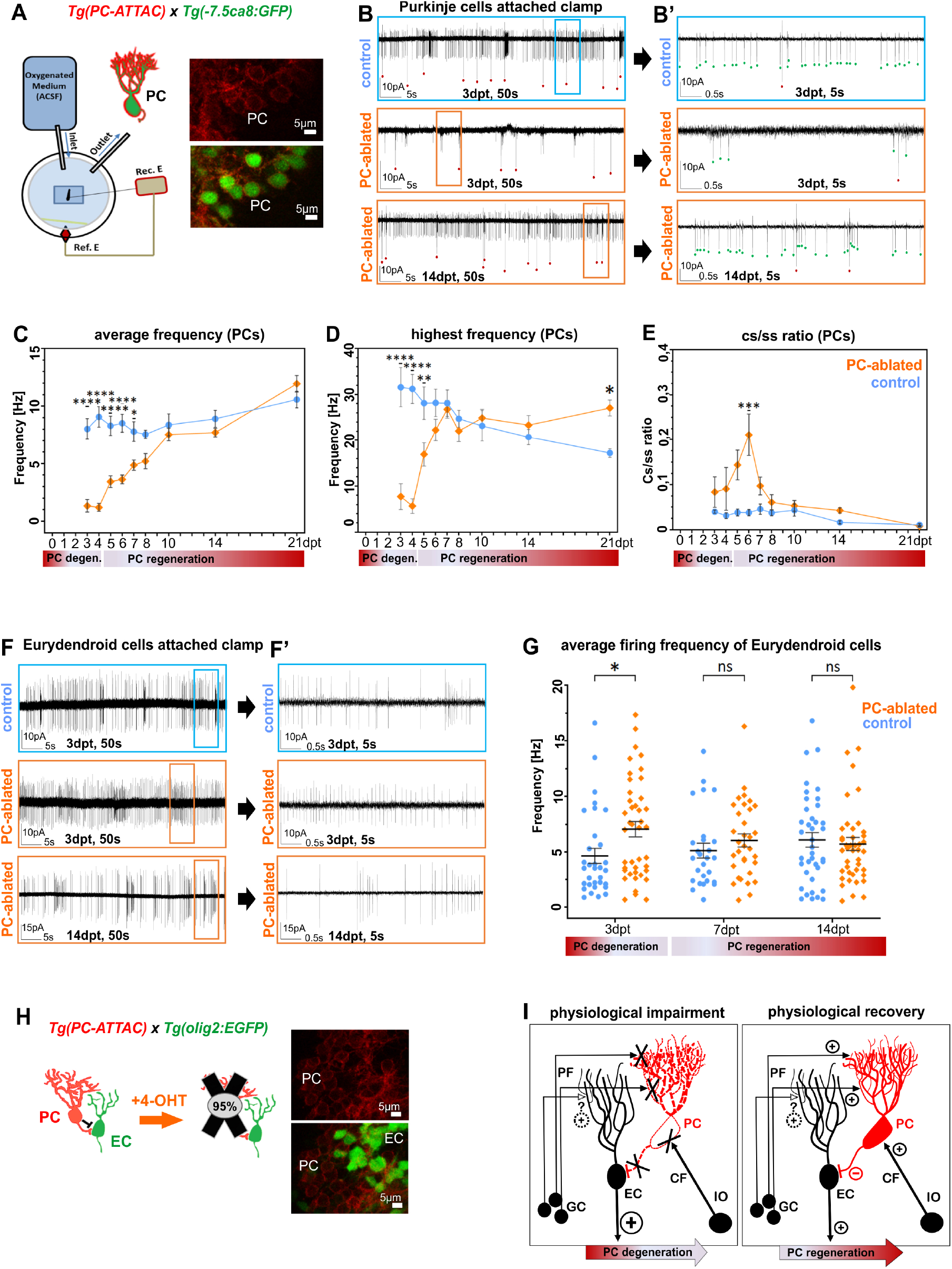
Electrophysiological properties of PCs during ablation and regeneration phase. **(A)** Patch-clamp recording set-up used for all experiments. Fluorescent PCs in larvae of the double transgenic line Tg(PC-ATTAC)/Tg(−7.5*ca8*:EGFP). **(B)** 50s trace of representative recordings of the tonic firing activity in control larvae 3 days after EtOH or in PC ablated larvae 3 and 14 days after 4-OHT treatment and (**B’**) 5s traces for all 3 traces shown. Red dots mark complex spikes and green dots mark simple spikes. **(C-E)** Diagrams representing results of the electrophysiologic investigations in PCs. C) Average tonic firing frequency plotted vs dpt. D) Highest spontaneous bursting frequency over an interval of 1s during a 100s trace plotted vs dpt. E) Ratio of the complex spikes to simple spikes. **(F)** 50s trace of representative recordings of the tonic firing activity of ECs in control larvae 3 days after EtOH or in PC ablated larvae 3 and 14 days after 4-OHT treatment and (**F’**) 5s traces for all 3 traces shown. **(G)** Average tonic firing frequency of ECs plotted vs dpt. **(H)** Illustration of PC loss after 4-OHT treatment and representative image of the PC layer from a double transgenic Tg(PC-ATTAC)/Tg(*olig2*:GFP) larva. **(I)** Illustration of physiological impairment and recovery model of the input/output of PCs during PC degeneration and regeneration respectively. **Figure Supplement 2A-C.** Electrophysiological reference data of differentiating PCs. **Figure Supplement 2D.** Quantification of PC ablation efficiency in specimens for electrophysiological recordings. **Figure Supplement 2E.** Quantification of EC ablation efficiency in specimens for electrophysiological recordings. **Figure Supplement 2F-H.** Electrophysiological data of regenerating PCs. **Source Data 1.** Average firing frequency determination of PCs for Figure 3C, highest burst frequency numbers of PCs for Figure 3D, dataset of complex spike to simple spike firing ratios of PCs for Figure 3E. Average firing frequency determination of ECs for Figure 3G.

Next, reappearing PCs after ablation in PC-ATTAC^TM^ larvae in the Tg(−7.5*ca8*:GFP)^bz12^ background with ablation rates above 90% were investigated (**Figure 3A-E - figure supplement 2D, F-H)**. During the acute phase of PC degeneration and debris clearance (0-4 dpt), the average frequency and the bursting frequency of few remaining PCs in PC-ablated larvae were significantly lower compared to controls and also the cs/ss ratio was elevated in an obvious trend (**Figure 3C-E - figure supplement 2F-H**). During the acute regeneration phase (5-10dpt) increasing numbers of PCs displayed characteristic simple and complex spike patterns (Fig. 3B, B’) with the average tonic firing and spontaneous burst frequencies steadily increasing to normal mature PC values. Similarly, the cs/ss ratio after an initial increase, reestablished ratios of mature PCs. During the extended regeneration period (10-21dpt) the average tonic firing frequency and cs/ss ratio remained stable and compared well to mature control PCs. This indicates that regenerating PCs successfully integrated into the remaining cerebellar network within 10 days. Of note, the average spontaneous burst frequency of regenerated PCs was increased by 1,5-fold in regenerated PCs compared to controls at 21dpt (Figure 3D) suggestive for a possible compensatory mechanism (see discussion).

Eurydendroid cells (ECs) are the principal direct efferences of zebrafish PCs and represent the neuronal equivalent of deep cerebellar nuclei neurons in mammals (Heap et al., 2013; Matsui et al., 2014). PC-ATTAC^TM^ carriers were crossed into the Tg(*olig2*:EGFP)^vu12^ background, in which ECs are marked by EGFP expression (Figure 3H) (McFarland et al., 2008), to analyze consequences of PC ablation and regeneration for EC physiology. As expected, at 3dpt the average firing frequency of ECs was significantly elevated upon acute loss of PC inhibitory input (**Figure 3F, F’, G - figure supplement 2E**), as ECs may still receive excitatory input from parallel fibers (Hashimoto and Hibi, 2012; Hibi and Shimizu, 2012). This elevated firing frequency returned already at 7dpt to a slight non-significant elevation during ongoing PC regeneration, and was indistinguishable from ECs in controls at 14dpt (**Figure 3G - figure supplement 2E**). This suggests that regenerating PCs quickly reestablish proper inhibitory input with their direct efferences (Figure 3I). Alternatively, ECs adapt in their intrinsic firing properties to the lack of inhibitory PC input to compensate for an excessive parallel fiber excitation. Further mechanistic studies are needed once further neuroanatomical evidence for likely direct synaptic contacts between parallel fiber and ECs have been provided.

### Functional recovery by larval PC regeneration

To address visuo-motor behavior, PC-ATTAC^TM^ larvae with a PC-ablation efficiency of above 90% (**Figure 4 - figure supplement 1H**) were mounted in agarose to evaluate cerebellum dependent optokinetic response (OKR) performance (Kazuhiko Namikawa et al., 2019; Ying-Yu Huang, 2008). Right eye rotations induced by horizontally moving stripes and fast return of the eye (saccades) were recorded as deflection angle with respect to body axis (Figure 4A, B). At 2dpt, when PC degeneration is prominent, the speed of eye movement during object tracking and saccade frequency were reduced compared to controls (Figure 4C **left panel, D**). Strikingly, at 10dpt when on average 29% of PCs had been reestablished (**Figure 4 - figure supplement 1H**), OKR response had recovered neither differing in eye movement speed nor in saccade frequency compared to controls (Figure 4C **right panel, D**).

**Figure 4:**
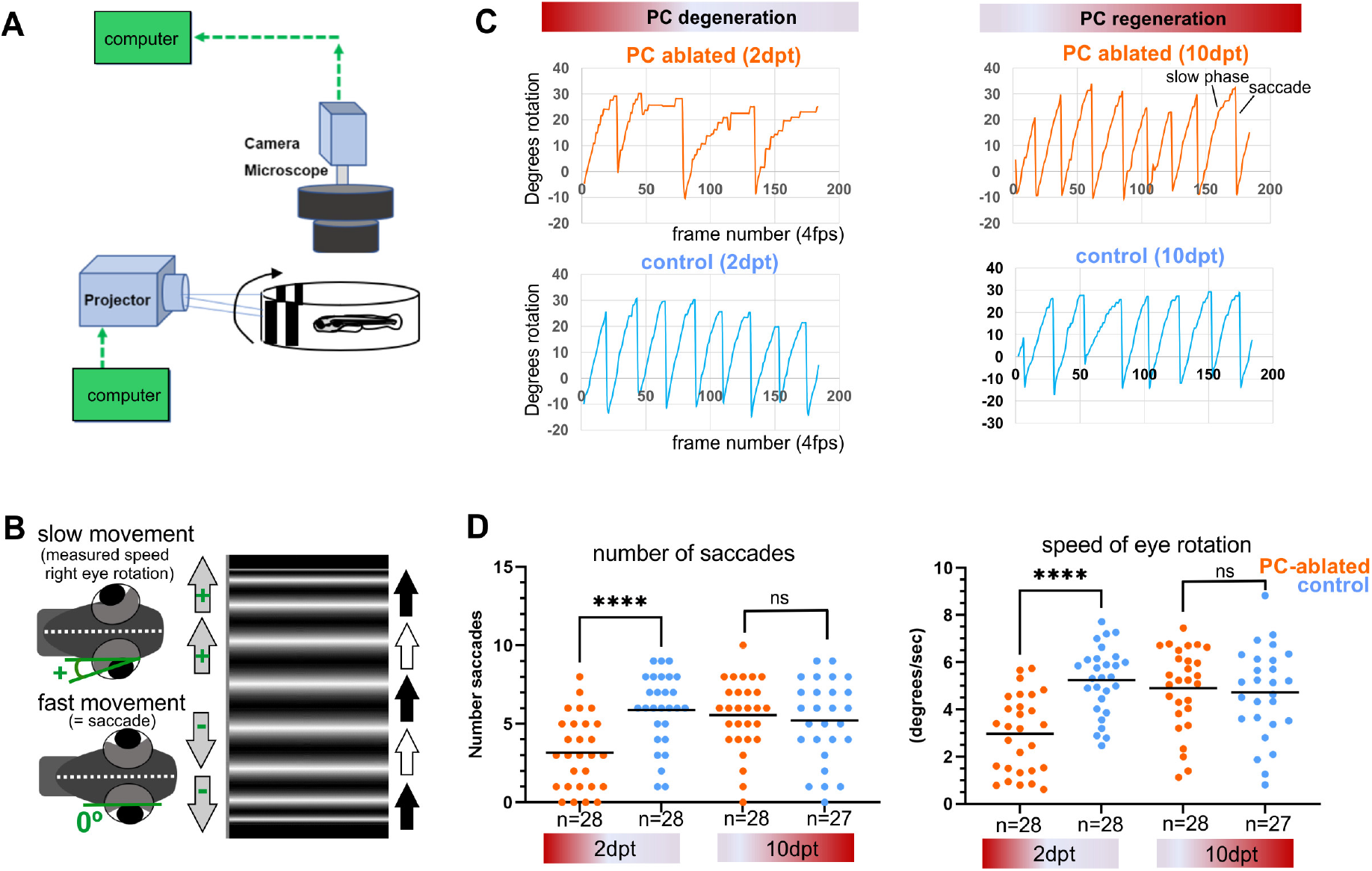
Visuo-motor behavior analysis: optokinetic response (OKR) after induced PC ablation in larvae. **(A, B)** Illustration of OKR set up (A) and eye movements in larvae (B) during performance of the OKR test. **(C)** Representative graphs showing OKR response performance during acute PC degeneration (2dpt) and regeneration (10 dpt) phases in PC-ablated (4-OHT treated) vs control group (EtOH treated). **(D)** Quantification of OKR response: number of normal saccades (eye rotation >19,5°) and speed of eye rotation during slow phase movements at 2 and 10 dpt. The data correspond to the results of 2 independent trials that were pooled. **Figure Supplement 1H.** Quantification of PC ablation efficiency in specimens for OKR testing. **Source data 1 and 2.** Datasets of eye deflection measurements during OKR behavior for Figure 4C, D. **Source data 3.** Eye rotation and saccade quantification for Figure 4C, D.

In order to evaluate cerebellum dependent locomotor behavior, swimming of individual control or PC-ablated PC-ATTAC^TM^ larvae were tracked in a 12-well plate for 6 minutes intervals (Figure 5A). Subsequently, PC ablation efficiency by counting PCs was determined to range above 90% (**Figure 5 - figure supplement 1I**). As EtOH treatment (the solvent for 4-OHT) can cause hyperactivity, while 4-OHT treatment is known to suppress hyperactivity (Blaser et al., 2010; Hoffman et al., 2016), respective solvent and compound controls were included. Also untreated and 4-OHT treated wild type larvae (to exclude effects from PC-ATTAC transgene expression) were analyzed, with the latter considered the most appropriate control. At 2dpt PC-ablated larvae traveled longer distances at elevated mean and maximum swim speeds suggesting hyperactive behavior, even above EtOH-treated controls (Figure 5D **upper row, E)**. Between 9-11dpt, when PC replenishment ranged on average around 40% (**Figure 5 - figure supplement 1I**), all swim parameters had recovered and were not different to all control groups respectively (Figure 5D **lower row)**.

**Figure 5:**
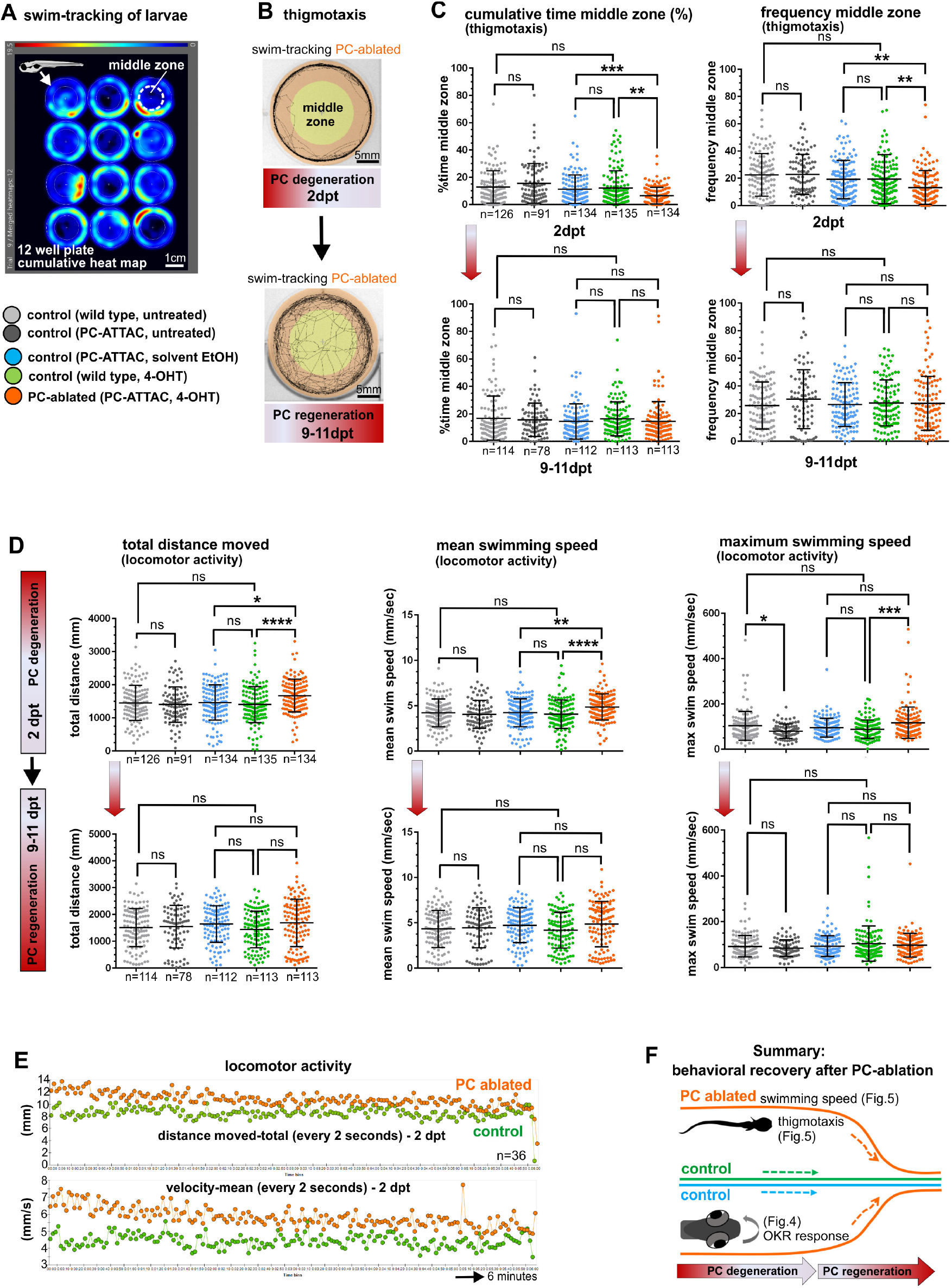
Swimming behavior analysis after induced PC ablation in larvae. **(A)** Heat map representing the location of zebrafish larvae in a 12 well plate during 6 minutes of swimming. **(B)** Example of swim track after PC ablation (2dpt) and during regeneration (10dpt) phases. **(C)** Quantitative analysis of swim preferences along the edge vs center zone of the arena (frequency of visits and percentage of time spent in the center zone) in PC ablated larvae vs control groups. **(D)** Quantitative analysis of locomotor activity (total distance traveled, mean and maximum swimming speed) in PC ablated larvae vs controls. **(E)** Graphs showing distance traveled and swim speed every 2 seconds during the tracking period, comparing PC ablated vs control larvae. **(F)** Illustration summarizing impairment and recovery of OKR (Figure 4) and locomotor behavior during PC degeneration and early regeneration phases respectively. The data from free swim tests correspond to the results from 4 independent ablations that were pooled. **Figure Supplement 1I.** Quantification of PC ablation efficiency in specimens for free-swimming analysis. **Source data 1.** Datasets of thigmotaxis and swim speed analysis for Figure 5C, D.

To analyze socio-emotional behavior, swim patterns in the above experiments were reevaluated for thigmotaxis. At 2dpt PC-ablated specimens spent significantly more time swimming along the walls and less frequently entered the open zone of the arena (Figures 5B, C **upper row**). Such an increase in thigmotaxis is considered to relate to elevated anxiety (Richendrfer et al., 2012). Again, at 9-11dpt PC-ablated specimens displayed no significant differences in thigmotaxis compared to all control groups (Figure 5B, C **lower row**).

Together these findings from behavioral analysis show that PCs not only reestablish the proper physiological signature after regeneration, but also regain their functional properties within 10 days in controlling visuo-motor, locomotor and thigmotactic functions already at a PC recovery rate close to 40% (Figure 5F).

### Recovery of adult PCs after induced ablation

The currently held view is that zebrafish PCs are only able to regenerate during larval stages, while this ability ceases in juveniles. Adult sexually mature zebrafish older than 3mpf are considered unable to replace lost PCs, like the mammalian cerebellum (Kaslin et al., 2017). The efficient functional regeneration of ablated PCs during larval stages lasting until adulthood prompted us to challenge this current view of limited cerebellar plasticity.

We therefore induced apoptotic cell death in heterozygous PC-ATTAC^TM^ adults at 5 months of age (Figure 6A) by three consecutive treatments with Endoxifen, an active metabolite of 4-OHT, which resulted in higher PC ablation efficiencies in adults compared to 4-OHT. As apoptosis of PCs and clearance of debris took longer than in larvae, remaining PCs were quantified on consecutive vibratome sections of the corpus cerebelli (CCe) at 13-14dpt (Figure 6C, F, I), which revealed an ablation efficiency above 90% compared to solvent controls (Figure 7A, B). Interestingly, while in rostral parts of the CCe PC ablation was nearly complete, PCs in the caudal part were more resistant to apoptotic ablation (Figure 6A, C, F).

**Figure 6:**
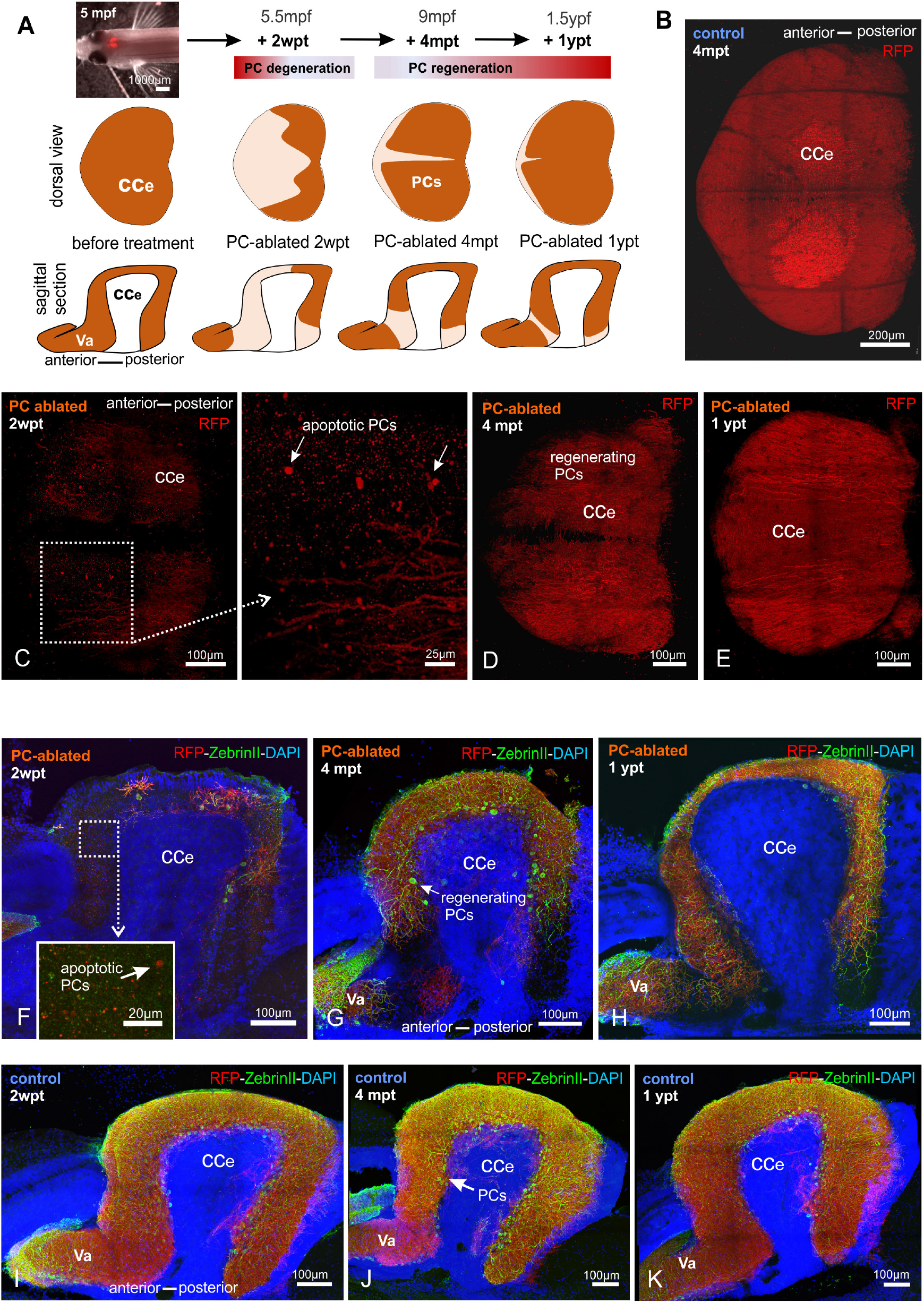
Monitoring of the PC layer after induced PC ablation in adults. **(A)** *In vivo* stereomicroscopy showing tagRFP-T fluorescence in the PC layer, and illustration of the time course of fluorescence recovery after PC ablation in adults (at 5mpf) monitored until 1 ypt. **(B-K)** Representative confocal images of whole mount cerebelli from dorsal view (B-E) and sagittal vibratome sections after immunostaining with the antibodies anti-tagRFP and anti-ZebrinII (F-K), comparing the cerebellum in ablated fish (endoxifen treated; C-E, F-H) vs control group (DMSO treated; B, I-K). Arrows in C, F point out apoptotic bodies. Equivalent results were observed in 2 additional independent ablations in adult cerebelli (data not shown). **Figure Supplement 3A-B.** RFP-intensity analysis in of adult regenerating PC layer **Figure Supplement 3C.** Quantification of size of PC layer based on PC fluorescence. **Figure Supplement 4A-C.** Distribution of *ptf1a*:GFP cells in adult cerebellum. **Figure Supplement 4D-I.** De- and regeneration of cerebello-vestibular tract in adult PC-ablated zebrafish compared to controls.

**Figure 7:**
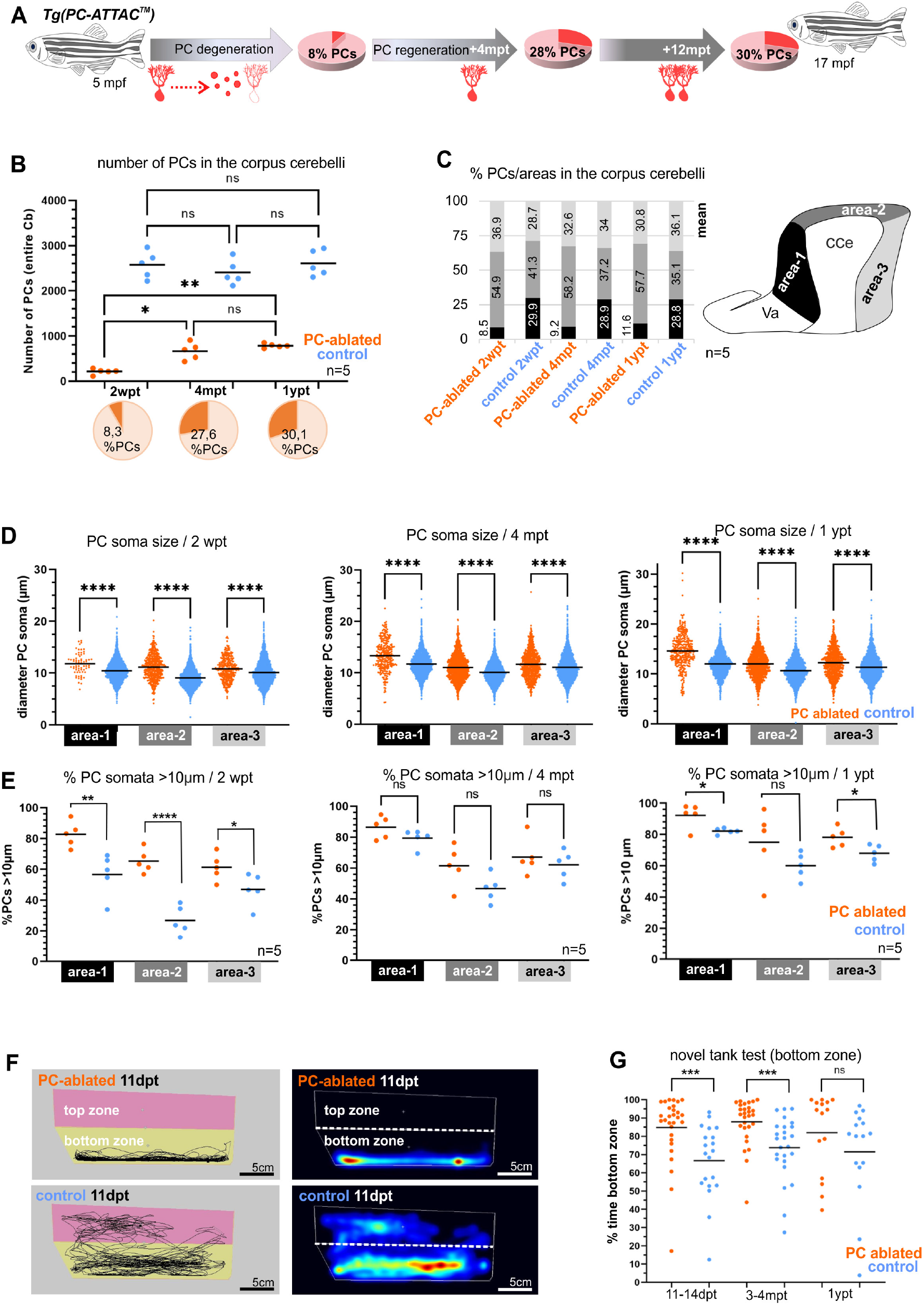
Quantitative analysis of PCs and behavior test during degeneration and recovery after induced ablation in adults. **(A)** Illustration of the percentage of PC layer replenishment during degeneration and regeneration phases. **(B)** Number and percentage of PCs after induced PC ablation monitored until 1ypt. **(C)** Subdivision of the CCe into rostral, dorsal and caudal areas (areas 1-3), and the respective proportion of PCs in each area. **(D)** Quantification of the diameter of PC somata in the different areas of the CCe. **(E)** Percentage of PCs with somata larger than 10µm from the total amount of PCs per area per brain. **(F)** Representative images of swim tracks and heat map of location of adult zebrafish in the novel tank test during 6 minutes. **(G)** Percentage of time spent in the bottom zone. The data correspond to the results from 3 independent PC ablations that were pooled. The cellular quantification in all graphs represent PCs from the entire CCe of 5 fishes per group per time point, as average from the whole PC population of each brain (B, C, E), or at single cell level that were pooled (D). **Figure Supplement 3D.** Quantification of body size of adult PC-ablated and control fish. **Figure Supplement 5A-C.** Quantification of regional distribution of regenerating PCs after adult PC ablation. **Figure Supplement 5D.** Quantification of PC soma size of regenerating PCs at different time points after PC ablation in adult zebrafish compared to controls. **Figure Supplement 5E.** Examples of different PC subtype morphologies in the adult cerebellum. **Figure Supplement 3E.** Quantification of swim speed and travelled distance of adult PC ablated zebrafish compared to controls during novel tank test. **Source data 1.** PC quantification for Figure 7A-C, PC somata measurements for Figure 7D, quantification of PC subtypes für Figure 7E, and exploration behavior in the novel tank test assays for Figure 7G. Detailed numbers and statistics shown in the figures are provided in Supplementary Table 1.

Fluorescent stereomicroscopy recordings from living fish confirmed the results of PC quantification from fixed tissue. At 2 weeks post treatment (wpt) control PC-ATTAC^TM^ fish displayed a strong and continuous red fluorescence from the dorsal surface of their CCe, while in PC-ablated specimens, fluorescence was faint at best, or completely absent (**Figure 6 - figure supplement 3A, B**). Strikingly, at 4 and 12mpt a robust cerebellum-derived fluorescence had reappeared covering an increasing area of the CCe (**Figure 6 - figure supplement 3C**). This fluorescence gradually converged to maximum intensity values obtained from adult controls (**Figure 6 - figure supplement 3A, B**), suggesting a significant PC regeneration. The observation of EGFP fluorescent cells in the CCe of Tg(*ptf1a*:EGFP) adult fish showed that a source of progenitor cells for PCs is still present at advanced adult ages (**Figure 6 - figure supplement 4A-C**).

PC quantification on sagittal sections by immunohistochemistry against TagRFP-T and the PC-specific epitope ZebrinII (Figure 6F-K) confirmed that 2 weeks after PC-ablation nearly any PC had survived in the adult CCe. Only few remaining PCs were preferentially located in caudal cerebellar regions (Figure 6F, I). At 4mpt ZebrinII expressing PCs with elaborate dendritic trees were distributed throughout a reappearing PC layer (Figure 6D, G, J) amounting to 28% of PCs found in controls (Figure 7A, B). Finally, at 12mpt *in vivo* fluorescent stereomicroscopy and immunohistochemistry revealed that gaps between PCs had disappeared and ZebrinII positive PCs were evenly distributed throughout a continuous PC layer (Figure 6E, H, K). Yet, the size of the PC population one year after ablation comprised to 30% in numbers compared to solvent controls (Figure 7A, B), which was not due to an overall growth reduction, as the average body length in both fish groups was nearly identical (**Figure 7- figure supplement 3D**). Of note, PC axon bundles projecting directly to the vestibular nuclei (Knogler et al., 2019; Kazuhiko Namikawa et al., 2019), which were completely degenerated at 2wpt, were reestablished at 4mpt and formed a prominent cerebello-vestibular tract at 12mpt (**Figure 6 - figure supplement 4D-I**).

These findings demonstrate that degeneration of the entire PC population in the adult zebrafish cerebellum does not represent an irreversible loss of these principal cerebellar neurons as previously thought. Instead, the adult zebrafish cerebellum maintains an impressive capability to regenerate significant numbers of PCs counteracting degeneration.

### Patterns of adult PC degeneration and regeneration

With an average of 8% PC survivors, the caudal area of the adult CCe was more resistant to induced apoptosis than the rostral area, in which PC survival was sparse. When PC regeneration in sagittal sections of the CCe was evaluated based on morphological landmarks, medial and caudal regions of the PC layer were more efficient in replenishing lost PCs compared to rostral areas, irrespective of subdividing the CCe into two or three areas (**Figure 7C - figure supplement 5A-C**). This patterned PC degeneration and regeneration may reflect the subdivision of the PC layer into different functional subcompartments along the rostro-caudal axis (Knogler et al., 2019; Matsui et al., 2014), and future elucidation of the underlying mechanisms promise to reveal important insights into PC maturation processes.

Recently, four different PC subtypes have been distinguished based on soma size correlating with a different dendrite morphology and physiology, but not being segregated with respect to their location in the PC layer (Chang et al., 2020) **Figure 7D, E - figure supplement 5E**). By measuring the size of PC somata all PC subtypes could be identified during de- and regeneration. Yet, compared to PCs in controls, apoptosis-resistant and regenerating PCs contained on average a larger soma size in all areas of the CCe, independent of subdividing the PC layer into two or three rostro-caudal compartments (**Figure 7D - figure supplement 5D**). This could be due to a higher physiological demand on the few surviving PCs. Alternatively, PCs of the largest subtype I with a soma diameter larger than 10µm are more resilient to apoptosis induction, and indeed in all rostro-caudal cerebellar compartments PCs with a large soma diameter (> 10µm) were significantly overrepresented at 2wpt, which was maintained but less pronounced during regeneration up to 1 year (Figure 7E). Noteworthy, the anterior CCe that is most sensitive to apoptotic PC ablation, displayed the largest percentage of PCs with a soma diameter above 10µm, ranging on average between 83% and 92% of the entire PC content (area-1, Figure 7E). These findings suggest that strong adapting subtype I PCs are more resilient to apoptotic cell death induction compared to PC subtypes II-IV.

### Functional recovery by adult PC regeneration

Functional PC regeneration implies the restoration of behavior caused by adult PC loss known to result in compromised exploration (Buchberger et al., 2021; Elsaey et al., 2021). In the novel tank test, adult zebrafish initially remain close to the bottom, but progressively start to explore the new environment (Egan et al., 2009; Fontana et al., 2021). Shortly after ablation despite the massive PC loss, zebrafish showed no significant difference neither in their mean swim speed nor in the total distance traveled compared to solvent treated siblings (**Figure 7F - figure supplement 3E**). This confirmed previous findings that zebrafish can largely relinquish PCs for locomotive control under non-strenuous conditions. In contrast, specimens with acute PC degeneration (2wpt) as well as during PC regeneration (4mpt) hardly ever entered the upper half of the new tank and displayed a significantly reduced exploratory behavior compared to controls (Figure 7F, G). Yet, in their home tanks, these fish fed successfully under continuous water flow. Importantly, after one year regeneration period this difference in exploratory behavior between PC ablated and control fish had disappeared, demonstrating in this context the functional recovery of the ablated PC layer (Figure 7G), despite the lower total PC content compared to age-matched controls (Figure 7A, B). As PCs in the rostral area of the CCe are associated with regulation of non-locomotor behavior such as socio-emotional behavior, these findings suggest that also the less efficient PC regeneration in the adult rostral CCe is sufficient to regain proper control over exploratory behavior.

## Discussion

Acute invasive injuries have demonstrated the ability of the teleostean cerebellum including zebrafish to regenerate damaged neuronal structures by proliferation, migration and differentiation of neuronal progenitors (Kaslin et al., 2017; Zupanc and Zupanc, 2006). With respect to PCs, regeneration in larvae has been deduced from the reappearance of PCs in injured cerebellar tissue. The proper physiological maturation of PCs and their contribution to cerebellum-controlled behavior has not been validated though, posing the question if reappearing PCs in larvae can be considered as proper replacement for damaged PCs. Furthermore, acute injury studies consistently established that zebrafish Purkinje cells follow the human paradigm: while PCs in the developing and maturing cerebellum can still be replaced, the adult cerebellum has lost the ability to regenerate PCs. This change in regenerative potential has been explained with the exhaustion of a suitable PC progenitor cell type in the adult zebrafish cerebellum (Kaslin et al., 2017, 2013).

Acute injuries, which may represent only the minor cases of cerebellar damage, have a massive impact on the surrounding neural tissue and are either locally restricted, as in the case of stab wounds, or lead to the destruction of the entire cerebellar architecture, as in the case for hemisphere removal or blunt force damage (Fleisch et al., 2011; Hentig et al., 2021; Kaslin et al., 2017). More commonly instead is the death of specific neuronal cell types and in particular PCs in the case of neurodegenerative diseases, for which PC regeneration has not been addressed in zebrafish so far. Therefore, the generalized view that zebrafish are unable to regenerate adult PCs needed to be reinvestigated by cell type specific approaches and importantly non-invasive procedures.

We have readdressed this open question of PC regeneration with the help of the PC-ATTAC^TM^ model, in which PCs can be ablated specifically by inducing apoptosis and reappearing PCs can be monitored by fluorescent protein expression (Weber et al., 2016). Using this model, we here confirmed that after depleting nearly the entire larval PC population, PCs regenerate to their full extent. First regrowing PCs occurred within 5-7 days and were derived from their natural progenitors expressing the transcription factor *ptf1a*. Interestingly, this replacement of PCs did not occur by a rapid wave of significantly increased progenitor cell proliferation, but instead by a continuation of cerebellum growth accompanied by a slow but steady replenishment of PCs in number, until a PC population indistinguishable in size from non-PC ablated control fish was reached.

An explanation for the slow but steady regeneration and stability of the circuitry can be derived from our physiological studies. Attached cell patch-clamp recordings from PCs in non-ablated control specimens confirmed that zebrafish PCs mature physiologically quite rapidly within 2 days, and the entire PC population displays stable electrophysiological patterns already at 8 dpf, 5 days after the first PCs are born (Hsieh et al., 2014; Knogler et al., 2019). In PC ablated specimens, reappearing PCs with a mature physiological pattern could be observed as early as 4dpt, with only marginal electrophysiological differences to control PCs at 8dpt (**Figure 3 - figure supplement 2F-H)**. This fast reestablishment of PC physiological patterns allows for a slow rate of PC regeneration. Obviously, a proportion of the PC population (about half at 14dpt **Figure 3 - figure supplement 2D**) is able to recover the physiological properties of the entire PC population efficiently.

We showed that during the acute phase of PC apoptosis this loss of inhibitory input resulted in ECs, the direct PC efferences, in a significantly increased average firing frequency of 57% at 3dpt. With the reappearance of PCs at 4dpt, a reduction in EC firing frequency back to normal values at 7dpt was observed. This suggests that reappearing PCs quickly reintegrate into the PC circuitry. Thus, larval PCs do not only regenerate in cell number, but likely establish proper synaptic contacts. Alternatively, ECs adapt by plasticity to the lack of PC input along the same time-course.

Interestingly, the pattern of physiological maturation appeared to be the same between wildtype and regenerating PCs. An initially increased complex to simple spike ratio was reduced during PC maturation from 5 to 7dpf, similarly to firing patterns recorded from regenerating PCs from 5 to 7dpt. This alteration in complex to simple spike ratio has been suggested to be derived from multiple climbing fiber innervation of PCs that is subsequently reduced by pruning until the strongest climbing fiber input remains (Hsieh et al., 2014). Once proper genetic targeting of inferior olivary neurons is available, it will be an interesting question to compare climbing fiber innervation in wildtype and regenerating PCs with adaptations in electrophysiological dynamics.

A fast recovery of mature and stable physiological patterns in regenerating PCs predicts a similarly quick recovery of PC function in governing behavior. In zebrafish larvae, impaired PC function has been shown to result in eye movement defects by means of saccade performance during the optokinetic response (Matsui et al., 2014; Kazuhiko Namikawa et al., 2019), to lead to decreased exploratory behavior in anxiety-related tests and to cause delayed recovery from conditioned fear response (Elsaey et al., 2021; Koyama et al., 2021; Matsuda et al., 2017; Kazuhiko Namikawa et al., 2019). Indeed, we observed such behavioral deficits during acute PC apoptosis at 2dpt, but also the functional recovery for visuomotor behavior (optokinetic response), locomotor activity (hyperactivity) and socio-emotional behavior (thigmotaxis), between 9-11 days post-ablation, when the PC layer is only partially replenished (below 50% PC regeneration). These findings prove that PCs selectively ablated in larvae by apoptosis regenerate by repopulating the PC layer of the cerebellum in adequate numbers accompanied by the reestablishment of proper physiological and functional control. Thus, larval PC regeneration can be considered a true functional regeneration.

Studies on acute cerebellum injuries have established that PCs lose this ability to functionally regenerate during juvenile and young adult stages at about 3 months post-fertilization (Hentig et al., 2021; Kaslin et al., 2017). However, localized injuries could be counteracted by functional plasticity of remaining PCs nearby, which may be faster compared to neuronal replenishment, thereby eliminating the need for a regenerative response of PC progenitor cells. Large scale injuries instead may remove or damage the proper cellular environment for regeneration.

We have therefore reinvestigated the potential of PC regeneration in the adult zebrafish cerebellum in the context of PC specific cell ablation clearly beyond young adult stages starting at 5 months post-fertilization. This approach more closely reflects a degeneration of the PC population, as it occurs in the many known human ataxias. Surprisingly, we observed a strong and extensive regenerative response reestablishing about 28% of the entire PC population within 4 months and 30% within one year. Moreover, PCs also reestablished direct efferent projections to the vestibular nuclei in the ventral hindbrain. Compared to larvae, both the process of PC ablation and PC death as well as the regeneration required extended time periods. This is likely due to both a deceleration of neurogenesis and a reduced progenitor population (Kani et al., 2010; Kaslin et al., 2017) as well as a compact organization of a fully differentiated cerebellum with a larger size in adults compared to larvae.

Behavioral tests revealed that PC regeneration in adults was accompanied by recovery of PC-mediated compromised exploratory behavior. Therefore, PCs do not only reappear, but also reestablish characteristic functions of a wildtype PC population. Furthermore, these findings suggest that about one third of the size of the adult wildtype PC population is sufficient to reestablish cerebellum-controlled behavior maybe eliminating a need for further PC regeneration. Clearly, the adult zebrafish cerebellum is capable of significant functional PC regeneration under conditions of cell type specific PC loss.

A recently established zebrafish model with compromised adult PC functions displayed defects in both exploratory as well as locomotor behavior (Buchberger et al., 2021). Interestingly, between PC-ablated PC-ATTAC^TM^ and control specimens speed of swimming and distances travelled were not significantly different, but only exploratory behavior. This is consistent with findings in a recently generated zebrafish model of PC degeneration (Elsaey et al., 2021), and both PC degeneration models have in common that loss of PCs is non-homogenous with a more extensive PC loss in the anterior adult CCe and more resilient survivors in posterior regions. Intriguingly, such a patterned PC degeneration has been found in mouse models of several human cerebellar disorders in which anterior cerebellar regions are stronger affected by PC degeneration than posterior ones (Sarna and Hawkes, 2011, 2003). Moreover, regeneration rates differed similarly as PC replenishment occurred faster in posterior than in anterior cerebellar regions. These observations support the view that suggested a functional pattern in the zebrafish cerebellum with posterior regions of the CCe predominantly controlling locomotor functions, while anterior cerebellar regions are more involved in mediating non-locomotor behavior such as socio-emotional behavior (Chang et al., 2021b, 2020; Harmon et al., 2017; Knogler et al., 2019, 2017; Matsui et al., 2014).

In addition to regional differences, patterned PC death in mammalian cerebellar diseases is also dependent on PC subtypes (Sarna and Hawkes, 2003). The existence of different PC subtypes in the adult zebrafish cerebellum has recently been established based on physiological criteria correlating to differences in PC soma size (Chang et al., 2020). PCs with soma diameters larger than 10µm (PC type I) were significantly overrepresented in our studies after acute PC ablation and during adult PC regeneration. A simple explanation for this finding is that PCs in a depleted PC population use the available space for increasing in size. Alternatively, remnant PCs grow in size because of plasticity processes with PCs trying to recover as many functions as possible. Another explanation may be that the observed difference in PC soma distribution reflects a difference in resilience of PC subtypes to neurodegenerative processes or a different regenerative potential among PC subtypes. Addressing these questions by establishing the proper technical tools for investigating the regional as well as the subtype differences in PC regeneration represent future rewarding experimental goals to further reveal the mechanistical concepts and limitations of PC regeneration in the adult cerebellum.

## Acknowledgments

We gratefully acknowledge Kazuhiko Namikawa, Jakob von Trotha and Florian Hetsch for helpful advice and experimental support. We thank Marcus Semtner (Max Delbrück Center, Berlin) for providing us with access to the Igor code for the analysis of electrophysiological data, and Janine Fichtner for support in their graphical presentation. We thank Alexandra Wolf-Asseburg, Iris Linde and Timo Fritsch for excellent technical support, and all members of the Köster group for critical discussions. This work was supported by the European Union (Horizon 2020 research and innovation program under the Marie Sklodowska-Curie actions Individual Fellowships H2020-MSCA-IF-2015, grant agreement No 703961, to S.P.M.) and the Bundesministerium für Bildung und Forschung (BMBF; Era-Net NEURON II CIPRESS to JCM).

## Author contributions

Conceptualization, SPM, PS, BW, RWK; methodology, SPM, PS, BW; investigation, SPM, PS, data analysis, SPM, PS, BW; all authors contributed to writing the manuscript; funding acquisition, JCM, KA, RWK.

## Data availability

Detailed numbers for statistics shown in the figures are provided in Supplementary Table 1. All data generated or analyzed during this study are included in the manuscript and supporting files.

## METHODS and MATERIALS

A list of key resources and their availability is provided in Supplementary Table 2.

### Animal husbandry

All animals were raised and kept in the zebrafish facility in accordance with established practices adhering to the guidelines of the local government (Aleström et al., 2019; Westerfield, 2007). AB wild-type or *brass* pigmentation mutant zebrafish were used for matings as indicated for each experiment. No selection criteria were used to allocate zebrafish of both sexes to any experimental group. Embryos and larvae were maintained in zebrafish 30% Danieau rearing medium (100% Danieau: 58 mM NaCl, 0.7 mM KCl, 0.4 mM MgSO_4_, 0.6 mM Ca (NO_3_)_2_, and 5mM HEPES, pH7.2) at 28°C. Larvae older than 7 days and adults were maintained in fish tanks according to standard protocols at 28^°^C, under light-dark conditions (simulating day-night cycle), and constant running water exchange. Stable transgenic lines used in this work include the Tg(ca8:-FynTagRFP-T2A-Casp8-ERT2)^bz11^ line abbreviated (PC-ATTAC^TM^) (Weber et al., 2016) approved by LAVES of Lower Saxony, Germany (license AZ33.14-42502-04-068/07) and the reporter fish lines Tg(−7.*5ca8*:EGFP)^bz12^, Tg(*olig2*:EGFP)^vu12^, Tg(*ptf1a*:EGFP), and Tg(*gfap*:EGFP)^mi2001^. All experimental protocols for animal research were approved by governmental authorities of Lower Saxony, LAVES, (AZ33.19-42502-04-20/3593). All efforts were made to use only the minimum number of experimental animals necessary to obtain reliable scientific data.

### Purkinje cell ablation

For Purkinje cell ablation experiments in larvae (4 to 6 days post fertilization), 4-OHT (cis/trans-4-Hydroxytamoxifen, Sigma Aldrich, St. Louis, MO, USA) at a concentration of 8µM was added to the rearing medium for 18 to 20 hours. Since EtOH was used as solvent for 4-OHT, 0.4% EtOH was applied to the rearing medium of a control group for the same time period. On the next day, the medium was exchanged 3 times and the ablation rate of PCs was analyzed under a confocal laser scanning microscope (LSM). Ordinary ablation rates higher than 85-90% at day two post-treatment (dpt) compared to the control group were considered successful to use these larvae for further experiments.

To deplete PCs in adults, 5 months old zebrafish were incubated overnight in fish facility water supplemented with 4µM Endoxifen (Sigma Aldrich, St. Louis, MO, USA), an active metabolite of 4-OHT, which proved more efficient in adult PC ablations. Three consecutive treatments were performed each overnight at 28°C in the dark with resting and feeding periods during the day. Since DMSO is used as solvent for Endoxifen, control groups were treated with 0.13% DMSO. Afterwards, the chemicals were washed out at least 3 times with fresh fish facility water and specimens were returned to the fish facility.

### BrdU incorporation assay

For proliferation studies in larvae, 10mM BrdU diluted in Danieaús solution was added to the fish water after 4-OHT treatment in a 24 well plate (1ml/well) for 24 hours, or up to 5 consecutive days in case of cumulative BrdU analysis. After the treatment, larvae were euthanized and the brains were isolated and processed for immunohistochemistry. The number of proliferating cells in each group (4-OHT/EtOH solvent control) within the PC layer of the cerebellum, and therefore excluding the caudal upper rhombic lip, was counted using the cell counter plugin of the image analysis software FIJI (see below). Student t-test for two groups comparison (unpaired t-test two tailed) was applied.

### Brain tissue processing for immunohistochemistry and imaging

Brains of both, larvae and adults, were isolated and fixed by immersion in 4% paraformaldehyde (PFA) overnight at 4°C. For isolation of larval brains, fish were euthanized with an overdose of Tricaine and prefixed with PFA for 20-30 minutes, to allow for complete removal of the skin and brain isolation (using a minute pin 00 as dissection tool). After overnight fixation of the brain, PFA was washed out with an excess of phosphate buffered saline (PBS), and the tissue was processed to a methanol/PBS series (30, 50, 70, and 100%) for gradual dehydration and final storage in 100% methanol at −20^0^C.

### Immunohistochemistry

For immunostaining, larval brains were rehydrated in a methanol-PBS series (70, 50 and 30%) and washed several times in PBS. Larval brains were processed for immunostaining as whole mount. After washing with PBS, antigen retrieval with 10mM citrate buffer was achieved by heating samples in a water bath until boiling. After cooling to room temperature the brains were rinsed 5 times in PBS-T (PBS with 1% Triton X-100) for 5 min each, incubated in 100% precooled acetone for 15 minutes at −20°C to improve tissue permeabilization, followed by additional 5 PBS-T washing steps. Nonspecific protein binding sites were blocked in 5% normal goat serum in PBS-DT-I: PBS, 1% bovine serum albumin (BSA), 1% DMSO, 1% Triton X-100) for 1h at room temperature followed by primary antibody incubation in PBS-DT-II (PBS, 1% BSA, 1% DMSO, 0,3% Triton X-100) overnight at 4°C and additional 3hours at room temperature. The primary antibodies: Rabbit anti-tagRFP (1:1000, Evrogen AB233, Moscow, Russia), mouse anti-zebrinII (1:500, donated by Dr. Hawkes, University of Calgary, Canada), chicken anti-GFP (1:1000, Aves GFP-1010, Aves Labs, Davis, CA, USA) and rat anti-BrdU (1:200, Biozol GmbH 6326, Eching, Germany). Before anti-BrdU antibody application an additional incubation step with 2N HCl was performed for 30 min at 37°C to denature the DNA for better anti-BrdU antibody accessibility. Subsequently brains were washed 5 times in PBS-T several times for 5 min each at room temperature with constant agitation and incubated with the secondary antibody diluted in PBS-DT-II overnight at 4°C. Secondary antibodies: Alexa goat anti-rabbit 568 (1:500, Invitrogen A11011, Carlsbad, CA, USA), Alexa goat anti-mouse 488 (1:1000, Invitrogen A11001, Carlsbad, CA, USA), donkey anti-chicken-FITC (1:1000, Jackson ImmunoResearch 703-545-155, West Grove, PA, USA) and goat anti-rat 488 (1:1000, Invitrogen A-11006, Carlsbad, CA, USA). For DAPI (1µg/ml) stained brains were incubated for 30 min at RT followed by several PBS-T washes. Finally, brains were thoroughly rinsed in PBS-T and PBS before imaging.

With respect to adult brains, immunostaining was carried out on 70 µm brain vibratome sections (Leica VT1000S, Leica Biosystems GmbH, Nusslsoch, Germany). Brains were pre-embedded in 1% low melting agarose in PBS, and then embedded in 3% agarose. Vibratome settings: speed: 1mm/sec, vibration frequency: 75Hz. The immunostaining was performed as described above, except for the antigen retrieval and acetone incubation. All incubations steps using adult brain sections were performed on floating sections, and eventually transferred to glass slides for confocal microscopy imaging.

### Electrophysiological Recordings

For single cell patch-clamp recordings, zebrafish larvae were anaesthetized in a bath of **<u>z</u>**ebrafish **<u>a</u>**rtificial **<u>c</u>**erebro **<u>s</u>**pinal **f**luid (z-ACSF: 134 mM NaCl, 2.9 mM KCl, 2.1 mM CaCl_2_, 1.2 mM MgCl_2_, 10 mM HEPES and 10 mM d-Glucose at pH 7.8 [adjusted with 10 M NaOH]) containing 10 µM d-Tubocurare (Sigma Aldrich, St. Louis, MO, USA). After complete loss of motor functions, the medium was removed and the larva was embedded in 2% low-melting agarose diluted in z-ACSF. Skin and skull tissue were then carefully removed with a sharp thin glass needle without damaging the underlying brain tissue to expose the cerebellum (see (Schramm et al., 2021) for a detailed method description). The immobilized larva within a square agarose block was immediately glued to a coverslip and transferred to the recording chamber, which was constantly perfused with oxygenated z-ASCF. Prior to electrophysiological recordings, larvae were adapted to the chamber without any disturbance for 10 min to allow acclimatization and/or stress reduction after the skull surgery. For attached-cell recordings of Purkinje cells in “voltage-clamp mode”, electrodes with an inner resistance of 6-8 MΩ were pulled from borosilicate glass capillaries (Item#:1B150F-4, ID: 0,84 mm, OD: 1,5 mm, WPI Inc., Sarasota, FL, USA) on a micropipette puller (Model PC-10, Narishige Inc., Tokyo, Japan) and filled with z-ACSF. Purkinje cells were identified by cytosolic EGFP expression Tg(−7.5ca8:EGFP)^bz12^, (Kazuhiko Namikawa et al., 2019), and visualized with a fluorescence microscope (SCOPE-II, Scientifica Ltd., Uckfield, UK) equipped with a CCD camera (C10600, Hamamatsu, Photonics Europe GmbH, Herrsching, Germany). Patch-clamp electrodes were controlled by motorized micromanipulators (Scientifica Ltd., Uckfield, UK) for precise targeting of Purkinje cells. To reduce the risk of a pipette blockade, constant pressure was applied. Recorded signals were amplified with a TECH ITC-18 (HEKA Elektronik GmbH, Reutlingen, Germany). All Purkinje cells were clamped with a holding potential of 0 mV throughout the whole time of recording and in a loose patch configuration with seal resistance reaching between 40-200 MΩ. For every cell, all spontaneous events were recorded at a rate of 20 kHz for a time span of 100 seconds and then selected with a Gaussian filter at 1 kHz. All experiments were performed at room temperature (25°C). For reliable results and to guarantee vitality/ healthiness and normal cerebellar activity, larvae were only patched until a maximum of 1 hour after skull opening. The recorded cells were equally distributed between both hemispheres and regional localization. Also, 5 cells per fish were considered as maximum threshold for a fish and after the 5^th^ successfully patched/attached neuron the next larva was put in focus. At the end of the experiment, zebrafish were euthanized by immersion in 0.2% Tricaine.

For regeneration studies of Purkinje cells, double transgenic fish carrying the PC-ATTAC^TM^ and the −7.*5ca8*:EGFP transgenes were created by crossing carriers of the two transgenic lines Tg(*ca8*:FMA-TagRFP-2A-casp8-ERT2)^bz11^ and Tg(−7.5*ca8*:GFP)^bz12^. Double positive larvae were screened at 4 dpf and Purkinje cells were ablated by standard procedures. Double transgenic larvae incubated in 0,4% Ethanol for 20 hours served as controls. The ablation rate was monitored by imaging larvae from 0-2 dpt each day under a confocal laser scanning microscope (LSM) and cell number analysis was performed. If the ablation rate was stable among all larvae and at 2 dpt over 90% compared to controls of the same age, the experiment was considered successful and patch-clamp recordings and imaging was performed from 3-21 dpt (10-28 dpf).

For patch clamp recordings of eurydendroid cells fish of the reporter line Tg(*olig2*:EGFP)^vu12^ (Shin et al., 2003) were crossed with transgenic carriers of the Tg(PC-ATTAC^TM^)^bz11^ line. In the double transgenic offspring, ablation and cell number analysis were performed as described above. Eurydendroid cells were visualized by cytosolic GFP expression, driven by *olig2* regulatory elements. Patch clamp recordings were also performed in the attached cell configuration described above with electrodes from the same capillaries but with an inner resistance of 8-10 MΩ.

Electrophysiological data were recorded with the Patchmaster Software from HEKA (HEKA Elektronik GmbH, Reutlingen, Germany) and afterwards analyzed offline with Igor Pro 6.37 (Wavemetrics Inc., Portland, OR, USA) with a program code written by Marcus Semtner (Charité, Max Delbrück Center, Berlin, Germany). The counting of complex spikes was performed manually, whereas simple spike counting was automated. By this algorithm-based analysis all simple spikes were marked by the program, complete identification of all simple spikes was confirmed by subsequent visual inspection. Only traces which were stable for a minimum of 75 seconds and showed consistent spike patterns with comparable amplitudes were considered successful and taken for the final data analysis. The data sets from all recordings were collected in Excel 2019 (Microsoft Inc., Redmond, WA, USA) and then further analyzed and graphically represented with Prism9 (Graph Pad Software, San Diego, CA, USA). Statistical relevance was analyzed by two-way ANOVA with Šídák’s multiple comparisons test.

### Behavioral Experiments

#### Optokinetic response (OKR) in larvae

The set up for this visuo-motor behavior test consisted of a movie projector playing a stimulating movie with vertical black and white stripes moving horizontally in constant speed projected over 180° across a circular screen as previously reported (Matsui et al., 2014). Single fish were mounted in front of the screen, in the center of a 5 cm Petri dish. The eye movement response of the fish to the stimulus was recorded by a camera mounted to a stereomicroscope objective above the head of the specimens. For the quantification of OKR response, the rotation angle of the right eye with respect to the body axis was measured. A rotation in the direction of the stimulus corresponds to the slow movement phase (marked as positive degrees). The rotation in the opposite direction or return of the eye to the original position, corresponds to the fast movement phase or saccade (marked as negative degrees, rotations of more than 19.5° were counted as saccade). The larvae analyzed were individually embedded using 2% low melting agarose diluted in Danieaús solution. In order to analyze free movements of the eyes, a window in the agarose around the head was gently cut out, and the dish was filled with Danieaús solution (preventing distortion of the vision, as well as to ensure normal breathing and comforting conditions to the larvae). After 10 seconds of adaptation to the stimulus, the eye response was recorded for 45 seconds with a camera recording at 4 frames per second (independent trials without adaptation period showed consistent results). Afterwards, the larvae were released from the agarose, returned to the home tank, and finally sacrificed at the end of the experiment. The eye response was recorded (Leica Microsystems LasX software, Wetzlar, Germany) using a stereomicroscope (MZC3000G, Leica Microsystems, Wetzlar, Germany) that was equipped with a digital camera (DFC3000G, Leica Microsystems, Wetzlar, Germany). For more consistency of the test results, the OKR recordings were always carried out between 10am and 5pm with alternating test rounds of each group. The statistical test applied was unpaired Student’s t-test for parametric or normal distributed data.

#### Free swimming in larvae

The analysis of free-swimming using larvae was performed using a custom-built zebrafish box (Noldus Inc., Wageningen, The Netherlands), illuminated with infrared and regular bright field light. Specimens were automatically tracked with the Ethovision XT 12 software (Noldus Inc., Wageningen, The Netherlands). The larvae placed in a 15cm Petri dish were transferred to the behavior test room (at 28°C) for acclimatization at least 2 h before the experiment. Subsequently, larvae were placed in a 12 well plate (one larva per well; with approximately 3ml fish facility water/well) and let alone for an adaptation period of 5 minutes. The tracking was performed over a time period of 6 minutes. In order to quantify thigmotaxis, the arena was subdivided into middle and edge zones by the analytical software. Additional parameters measured were swim speed and total distance traveled. The larvae were returned to their respective fish tanks, and sacrificed at the end of the experiment. For more consistency of the results, the test was always carried out between 2:30pm and 5:30pm, with alternating of each group for swimming test analysis. The statistical tests applied were: ANOVA test for multiple group comparison (ordinary one-way ANOVA followed by the Šídák’s multiple comparisons test for parametric or normal distribution data or Kruskal-Wallis test for no-parametric data).

#### Novel tank test (NTT) in adults

The novel tank test was performed with a custom-built zebrafish box (Noldus Inc., Wageningen, The Netherlands) illuminated with infrared and regular bright field light, and automatic tracking with the Ethovision XT 12 software (Noldus Inc., Wageningen, The Netherlands). The fish were placed into behavior test room (at 28°C) approximately 1h prior to testing for acclimatization. For the novel tank test, the swimming of the fish was recorded for 6 minutes without a pre-adaptation period to the novel tank. The fish tank used for the test had a volume of 1,5 liter (Aquaneering Inc., San Diego, CA, USA). After finishing the tracking, adult fish were returned to their respective home tanks, and sacrificed at the end of the experiment. For the tracking, the arena was subdivided into upper and bottom zones by the analytical software. Due to some variability of the NTT behavioral assay, individual control fish that showed clear exploration behavior of at least 5% occupancy in the upper half of the tank were chosen for further analysis. Mann-Whitney two tailed test for no parametric data was applied.

### Imaging and data analysis

For imaging, larvae were anesthetized with 0.015-0.02% Tricaine (Sigma Aldrich, St. Louis, MO, USA) dissolved in 1% low melting agarose/Danieau solution. Images of the larval and adult zebrafish cerebellum were acquired using an SP8 laser scanning confocal microscope (Leica Microsystems, Wetzlar, Germany) with a 40x objective. Images were processed with the imaging and analysis Leica LasX software.

Additional analysis of data was carried out with following software: FIJI (for automated quantification from image data and quantification of the optokinetic response, freeware at https://imagej.net) and Ethovision (for swimming behavior quantification, Noldus Inc., Wageningen, The Netherlands). The quantification of PCs in juvenile and adult cerebelli was carried out after vibratome sectioning and immunostaining with anti-tagRFP and anti-ZebrinII antibodies. Semi-automated counting of cells was performed with the cell counter plugin of the FIJI software, in which every cell is initially identified manually followed by their automated detection with a z-stack of images to avoid, that cells are counted twice. The total amount of PCs per brain corresponds to the sum of all the sections. In order to minimize the error of double counting the same PCs from consecutive sections, cutting thick (70µm) vibratome sections was considered the most appropriate sectioning method.

The design of the figures was carried out with CorelDraw software (Corel Corporation, Ottawa, Ontario, Canada).

### Material Availability

Further information and requests for resources and reagents should be directed to and will be fulfilled by the Lead Contact.

## Supplementary Data

### Abbreviation list

4-OHT: 4-hydroxy-tamoxifen
CCe: corpus cerebelli
dpf: days post-fertilization
dpt: days post-treatment
Endox: endoxifen
EtOH: ethanol
fps: frames per second
GCL: granular cell layer
MOL: molecular cell layer
mpf: months post-fertilization
mpt: months post-treatment
PCL: Purkinje cell layer
PCs: Purkinje cells
Va: valvula cerebelli
VZ: ventricular zone
wpt: weeks post-treatment
ypf: years post-fertilization
ypt: years post-treatment

## Figure Legends

**Supplementary Figure 1:**
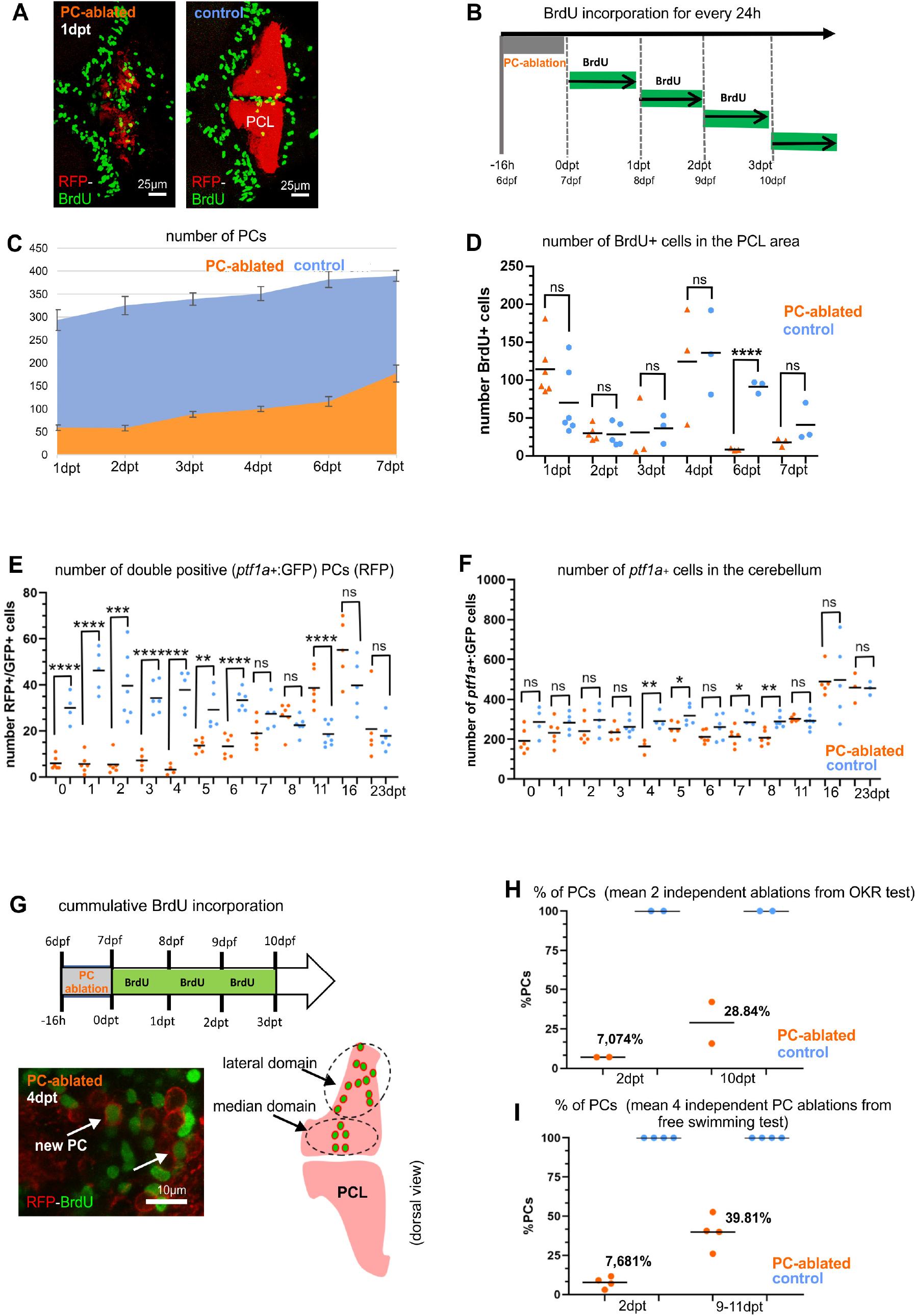
Cell proliferation and GABAergic cell progenitors in the cerebellum after PC ablation: BrdU analysis and average number of PCs after PC ablation in larvae, related to figures 1, 2, 4 and 5. **(A)** Representative images of BrdU positive cells in the cerebellum of PC ablated and control larvae. Rostral is to the left. (**B)** Schematic representation of the BrdU incorporation treatment and analysis carried out for periods of 24h. (**C, D)** Numbers of PCs (C) and BrdU positive cells within the PC layer area of the cerebellum -excluding the most caudal rhombic lip area-(D) during PC degeneration and beginning of regeneration phases. (**E)** Number of double positive PCs showing RFP expression in mature PCs, and green fluorescence from *ptf1a*-EGFP-expressing cells. **(F)** Number of *ptf1a*-expressing cells (GFP positive) in the cerebellum, a general increase of *ptf1a*-expressing cell progenitors after PC ablation is not detected. **(G)** Mature PCs with BrdU incorporation (after a cumulative treatment for several days), revealing the location of new PCs during the period of BrdU incorporation, grouped into median and lateral domains of newly generated PCs (this location is similar to newly generated PCs derived from *ptf1a*-EGFP-expressing progenitors). **(H, I)** Average percentage of PCs at acute degeneration phase and beginning of regeneration from independent trials of OKR and free-swimming tests after PC ablation in larvae. **Source data 1.** Quantification of BrdU-positive cells for Suppl. Figure 1D, quantification of *ptf1a*:GFP-expressing PC-ATTAC cells for Suppl. Figure 1E, quantification of *ptf1a*:GFP-expressing cells for Suppl. Figure 1F, quantification of PCs for Suppl. Figure 1H, I.

**Supplementary Figure 2:**
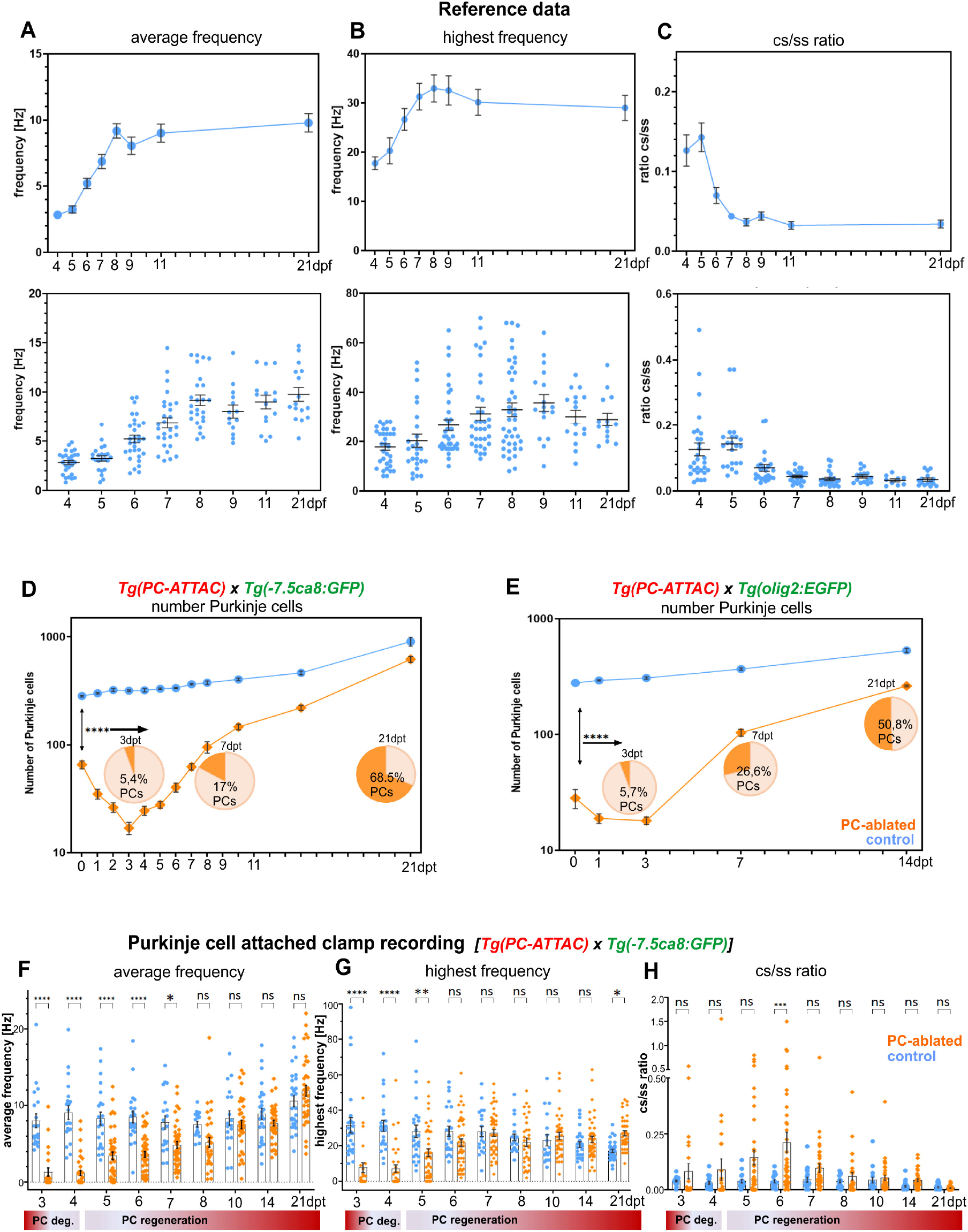
Electrophysiological reference data-set of larval Purkinje cell development and regeneration studies, related to figure 3. Figures **A-C** show the age-dependent physiologic activity of PCs, marked by green fluorescence in the reporter line Tg(−7.5*ca8*:GFP)^bz12^ under non-stimulated conditions over a time course between 4-21 days post fertilization. **(A)** line diagram of the average PC tonic firing frequency plotted against time over the entire time course, the corresponding single cell values are shown underneath in a box plot for each recording day. The two graphs in **(B)** show the highest firing frequency over a timespan of 1s during spontaneous burst events. The last two graphs in **(C)** show the time-dependency of the complex spike to simple spike ratio (cs/ss ratio). The line graph shows an overview of the population average over the entire time course, while the box plot illustrates the distribution of this ratio on the single cell level for each recording day. **(D)** PC number analysis during the electrophysiologic PC regeneration studies. **(E)** PC quantification during the electrophysiological studies of eurydendroid cells. The diagrams in **F-H** represent single cell results of PC activity during the electrophysiological investigations and support the graphs of Fig. 4C-E. **(F)** average tonic firing frequency plotted vs. days post treatment. **(G)** the highest spontaneous burst frequency over an interval of 1s during a 100s trace plotted vs. days post treatment. **(H)** complex spike to simple spike ratio (cs/ss ratio) of PC activity. Statistical significance in figures D-H was tested in a two-way ANOVA with Šídák’s multiple comparison test. **Source Data 1.** Average firing frequency determination of PCs for Suppl. Figure 2A, highest burst frequency numbers of PCs for Suppl. Figure 2B, dataset of complex spike to simple spike firing ratios of PCs for Suppl. Figure 2C. Quantification of PC numbers for Suppl. Figure 2D, E. Average firing frequency determination of PCs for Suppl. Figure 2F, highest burst frequency numbers of PCs for Suppl. Figure 2G, dataset of complex spike to simple spike firing ratios of PCs for Suppl. Figure 2H.

**Supplementary Figure 3:**
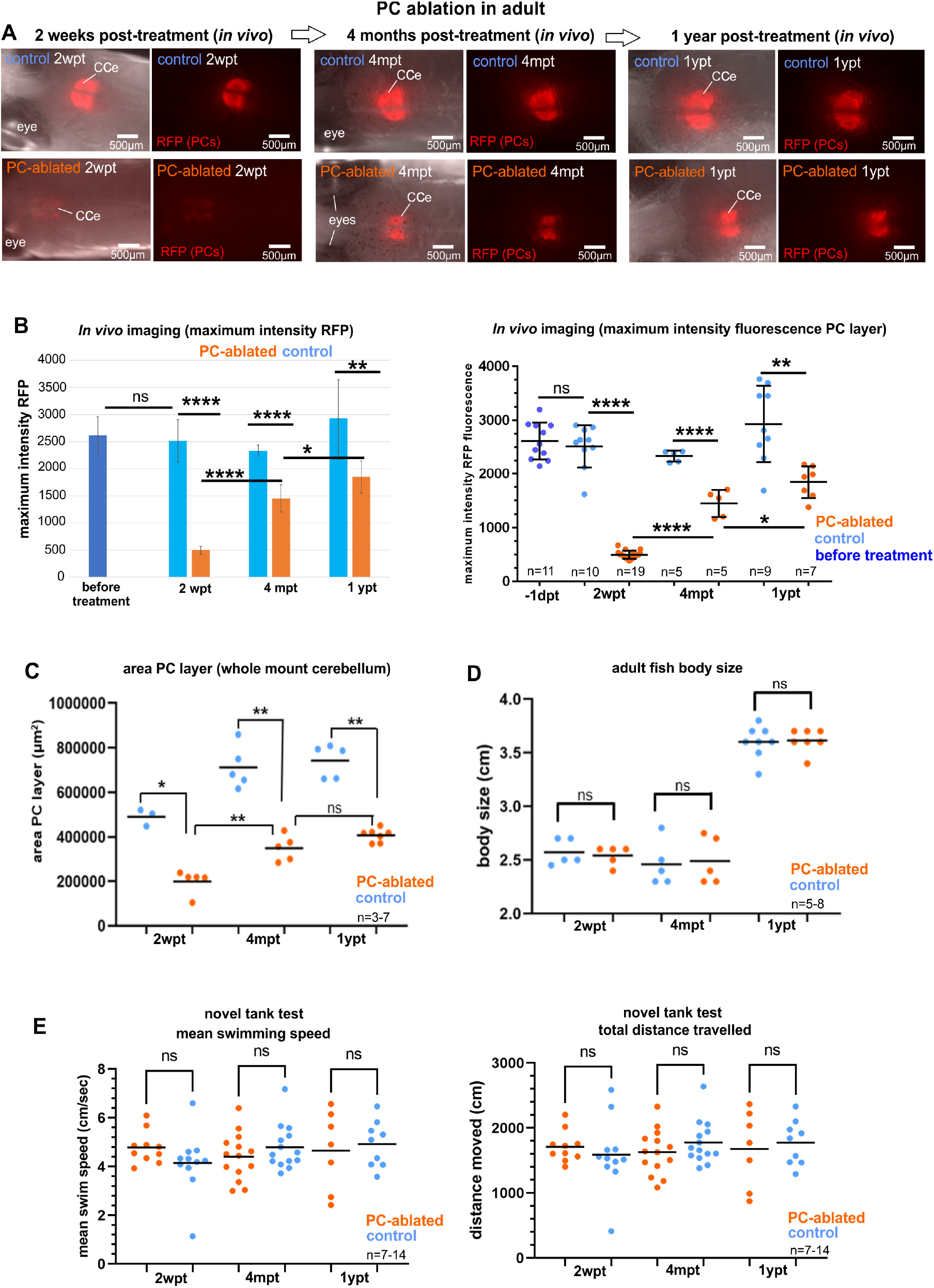
Imaging of the cerebellum *in vivo*, and quantification of the PC area, body length, and swimming after PC ablation in adult zebrafish; related to figures 6 and 7. **(A, B)** *In vivo* imaging of ablated PC layer in adults. Images recorded *in vivo* by stereomicroscopy revealing PC-derived RFP fluorescence in the PC layer, and measurement of fluorescence intensity. All fishes treated with Endoxifen showed very faint fluorescence in the cerebellum 2 weeks post-ablation, verifying actual PC ablation in the entire Endoxifen-treated group. The recovery of the PC layer displayed a significant increase of fluorescence at 4 months and 1 year post-ablation. (**C)** Area of the PC layer from whole mount confocal imaging during PC degeneration and recovery. The significant difference of the area compared to control fish displays the efficiency of PC induced apoptosis, and the regrowth of the area within the ablated group 4 months after the ablation shows the progressive regeneration of the PC population. (**D)** Body length of adult fish after induced PC ablation. No differences between fish from PC ablated and control groups were observed. **(E)** Novel tank test. Total distance traveled and mean swim speed after induced PC ablation in adults. No differences were observed between ablated and control fish during the PC degeneration nor during the regeneration periods were observed. For statistical analysis the Student t-test was applied for two groups comparison (unpaired t-test two tailed for parametric, or Mann-Whitney two tailed for no parametric data). **Source data 1.** Measure of maximum intensity of RFP fluorescence for Suppl. Figure 3B, area size determinations of PC layer for Suppl. Figure 3C, adult zebrafish body size measurements for Suppl. Figure 3D, mean swim speed and total distance values for behavioral analysis for Suppl. Figure 3E.

**Supplementary Figure 4:**
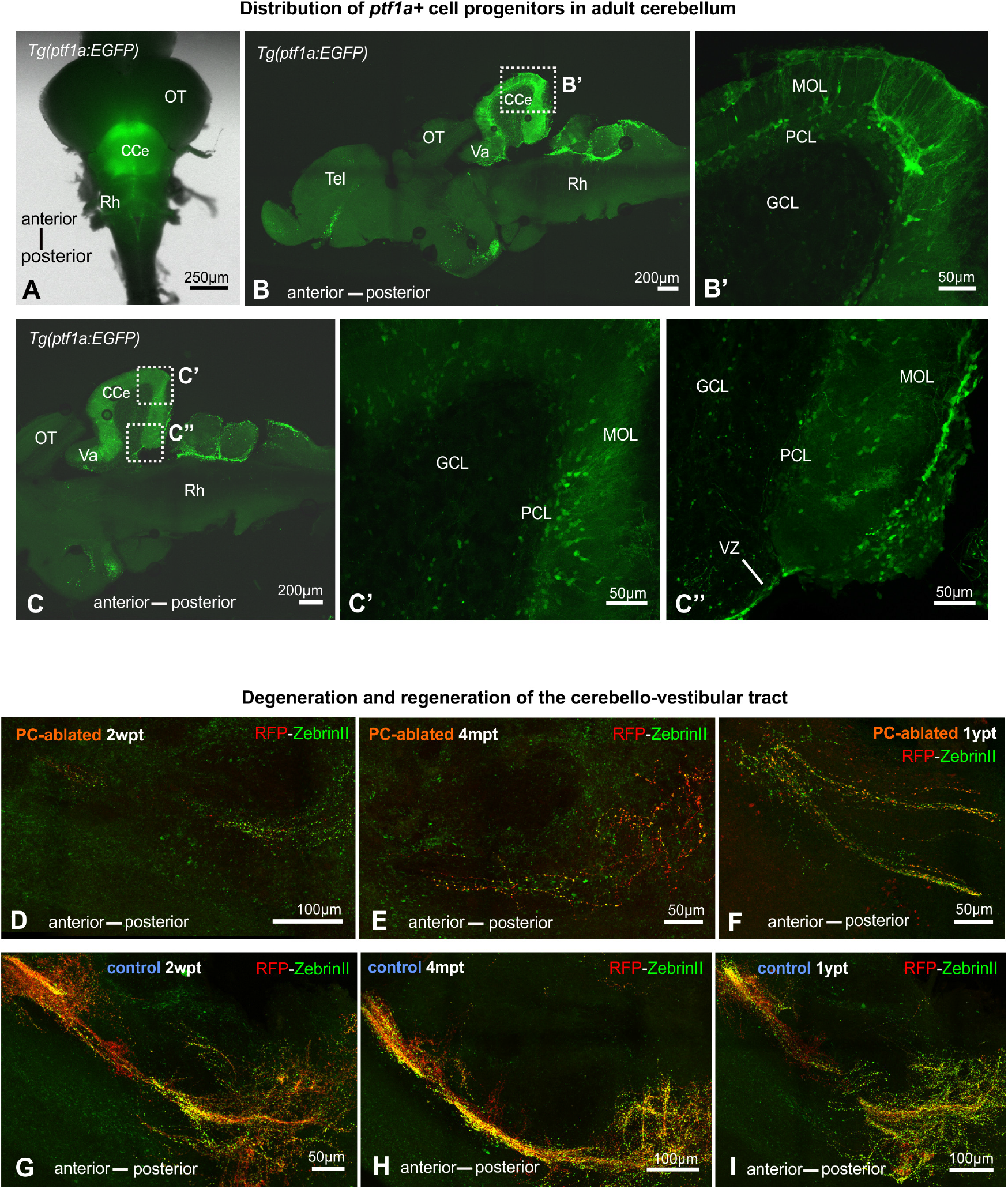
Distribution of *ptf1a*-expressing progenitors in the adult cerebellum (A-C) and degeneration and recovery of cerebello-vestibular tract after adult PC ablation (D-I), related to figures 6 and 7. **(A-C)** Representative images of whole mount brain (A) and sagittal sections (B, C) from the transgenic Tg(*ptf1a*:EGFP) reporter line of a 3 years adult still revealing the presence of *ptf1a*-expressing progenitor cells in the cerebellar PC and molecular layer in aged fish. **(D-I)** Images of the cerebello-vestibular tract after immunostaining with the antibodies anti-tagRFP and anti-ZebrinII on vibratome sections in PC-ablated fish (D-F) compared to control fish (G-I), show a vast tract degeneration at 2 weeks (D), and a partial recovery of this PC-derived axonal tract 4 months and 1 year after ablation (E, F). Rostral is to the left. Abbreviations: Cb cerebellum, CCe corpus cerebelli, GCL granular cell layer, MOL molecular cell layer, OT optic tectum, PCL Purkinje cell layer, Rh rhombencephalon, Tel telencephalon, VZ ventricular zone.

**Supplementary Figure 5:**
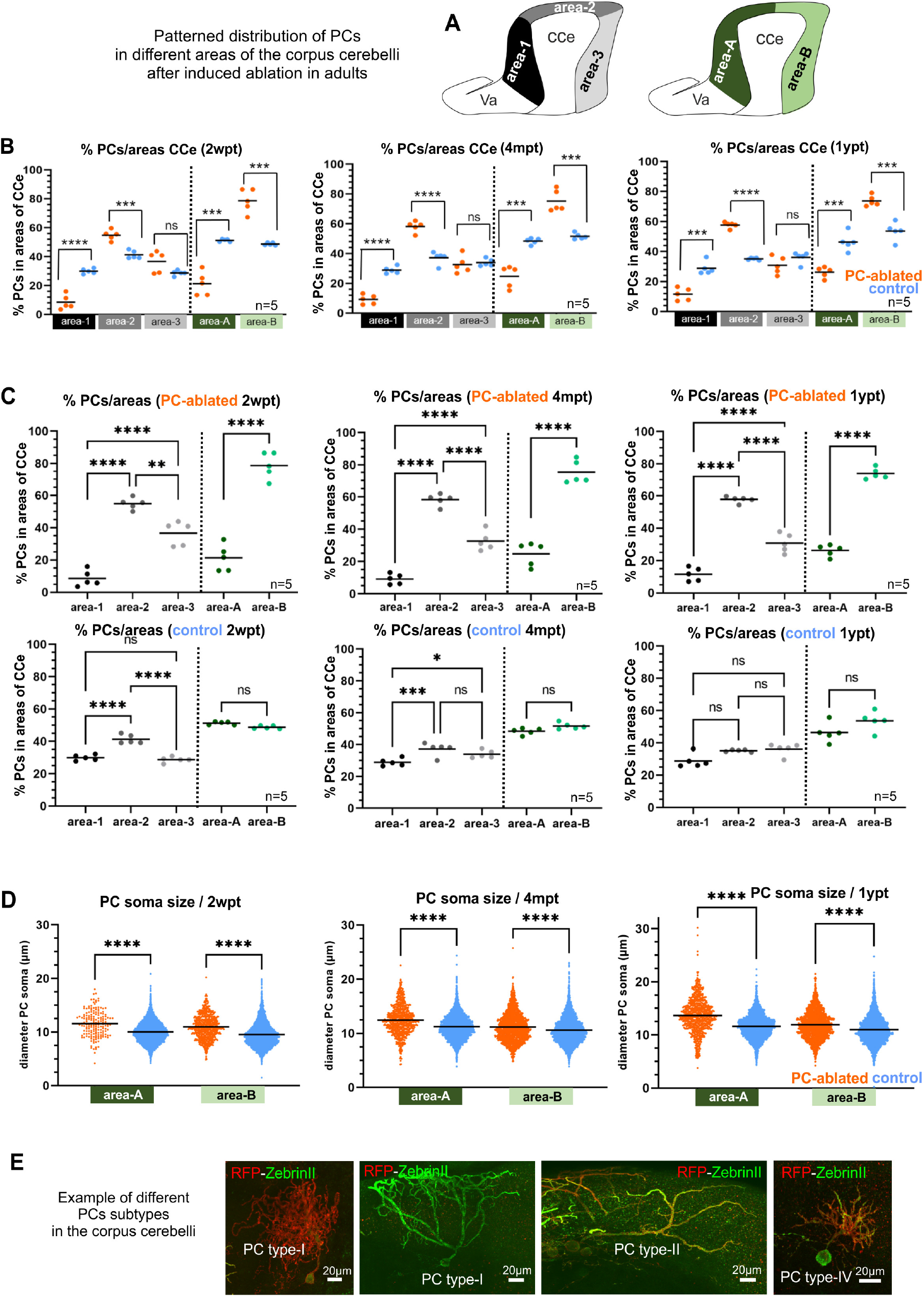
Quantification of patterned PC death and recovery throughout the antero-posterior axis of the CCe, related to figure 7. **(A)** Proposal of possible subdivisions of the CCe, into 3 areas (rostral = area 1, dorsal = area 2, and caudal = area 3) or into 2 halves (rostral = area A, and caudal = area B). **(B, C)** Percentage of PCs present in the different subdivisions of the CCe comparing ablated and control groups (B), and among fishes of the same group (C). Both groups, ablated and controls, present a tendency of lower amounts of PCs in the most rostral area than caudal parts of the CCe (C). However, when comparing between groups, the percentage of PCs in those rostral areas is significantly smaller in PC ablated fish than in control specimens (B). (**D)** Differences in the size of PC somata in the anterior and posterior halves of the CCe (areas A and B respectively). (**E)** Example images of different PC subtypes, visualized individually after induced PC apoptosis, which show different soma sizes and morphologies of their dendritic tree as described by (Chang et al., 2020). The quantification in all graphs represent PCs from the entire CCe of 5 fish per group per time point. Average values from the entire Purkinje cell population of each brain (B, C), or from individual PCs were displayed together (D). The statistical tests applied were: a) ANOVA test for multiple groups comparison (ordinary one-way ANOVA-followed by the multiple comparison Sidàk test for parametric or normal distributed data, or Kruskal-Wallis test for no parametric data), or b) Student t-test for two groups comparison (unpaired t-test two tailed for parametric, or Mann-Whitney two tailed for no parametric data). **Source data 1.** Distribution values of PCs for Suppl. Figure 5B, C. Measurements of PC diameters for Suppl. Figure 5 D.

**Supplementary Table-1.**
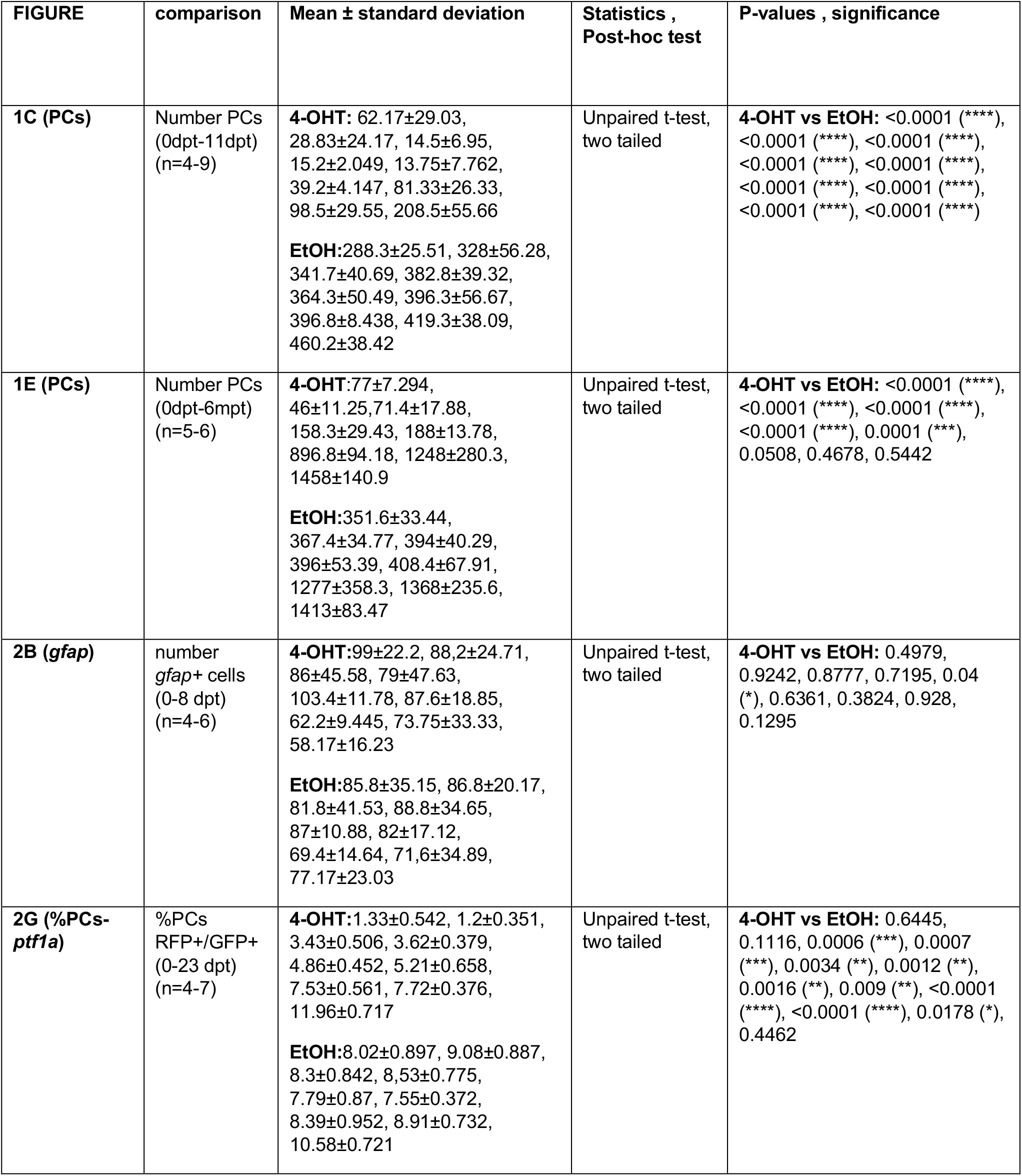
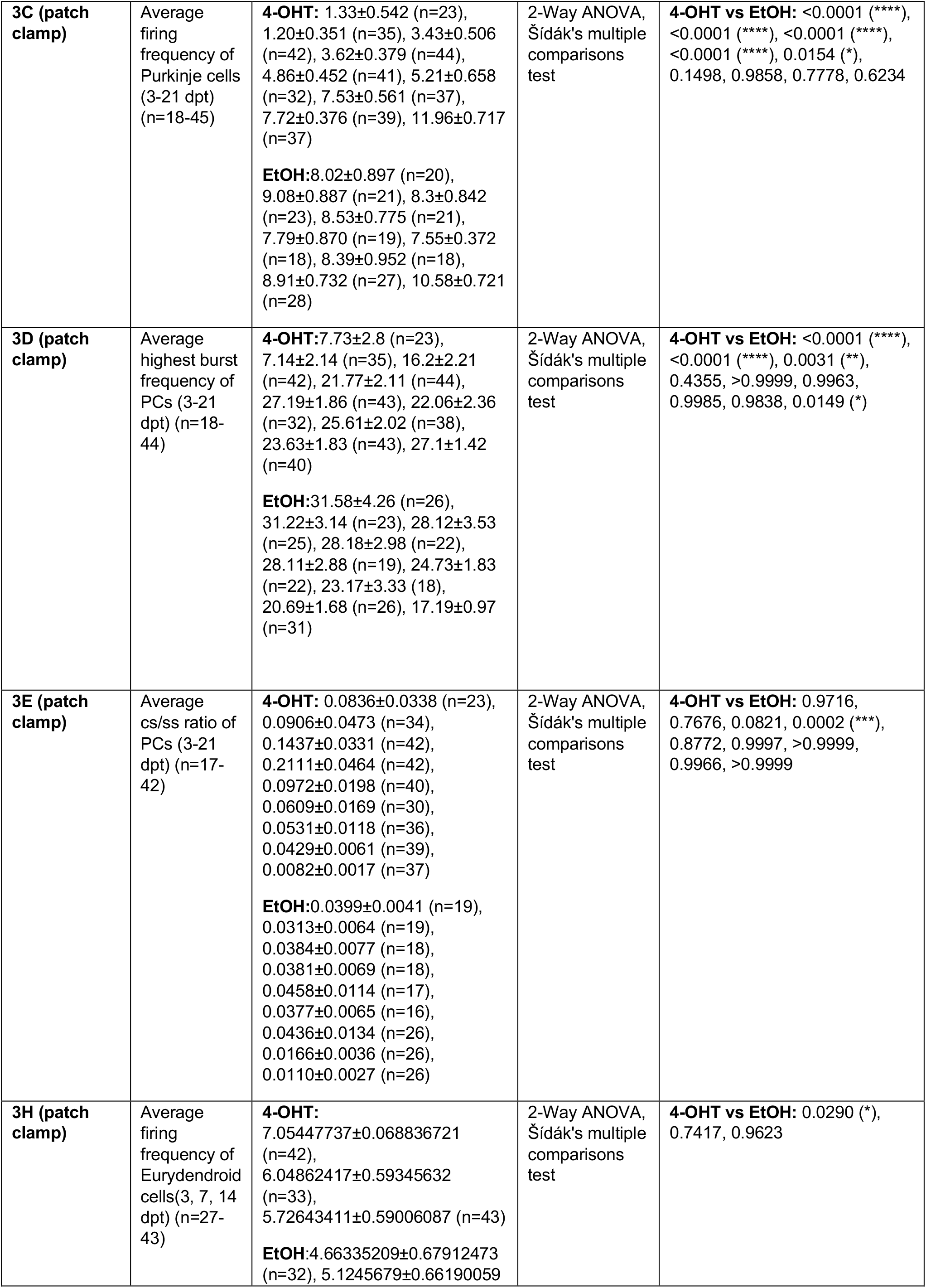
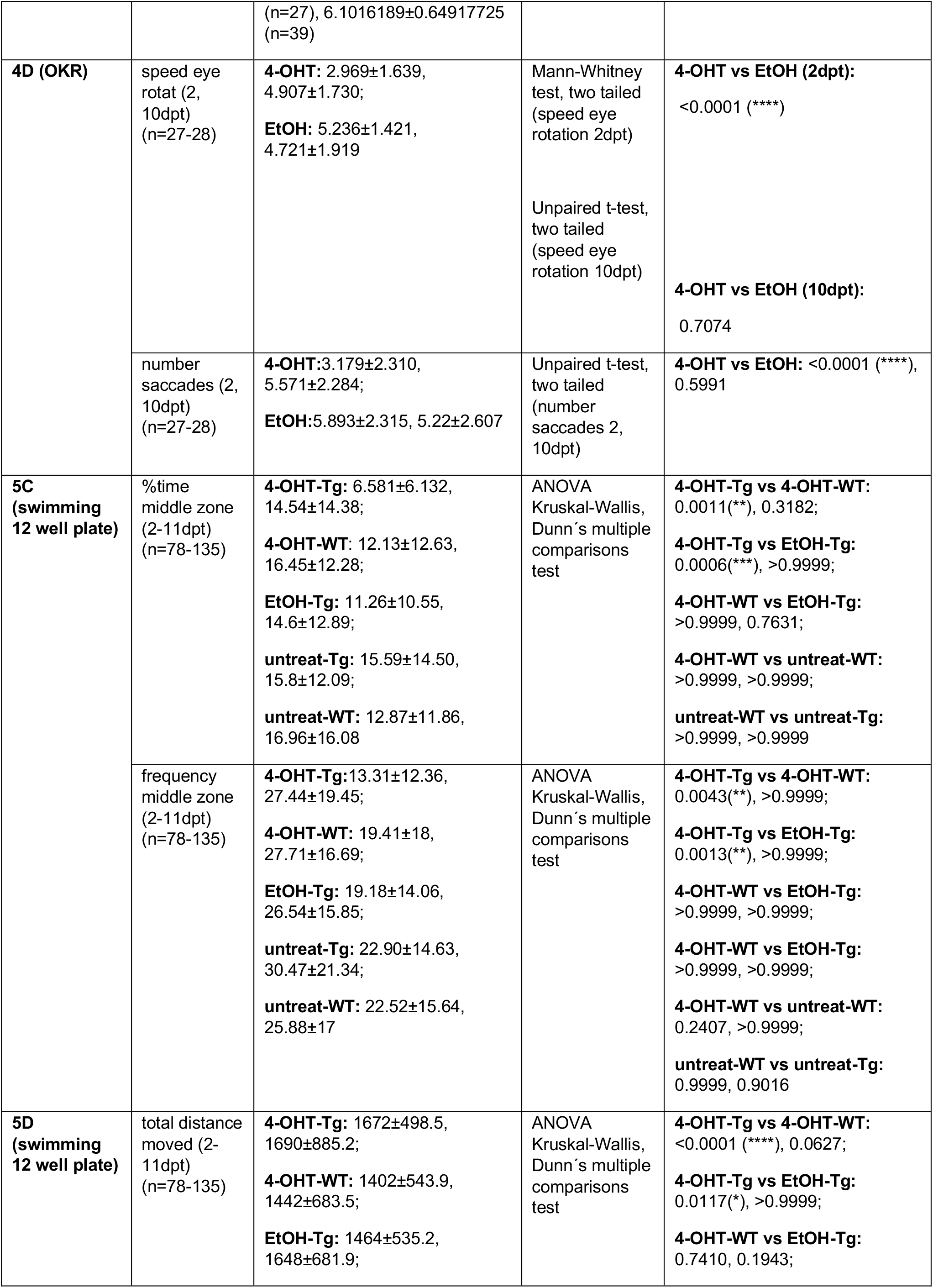
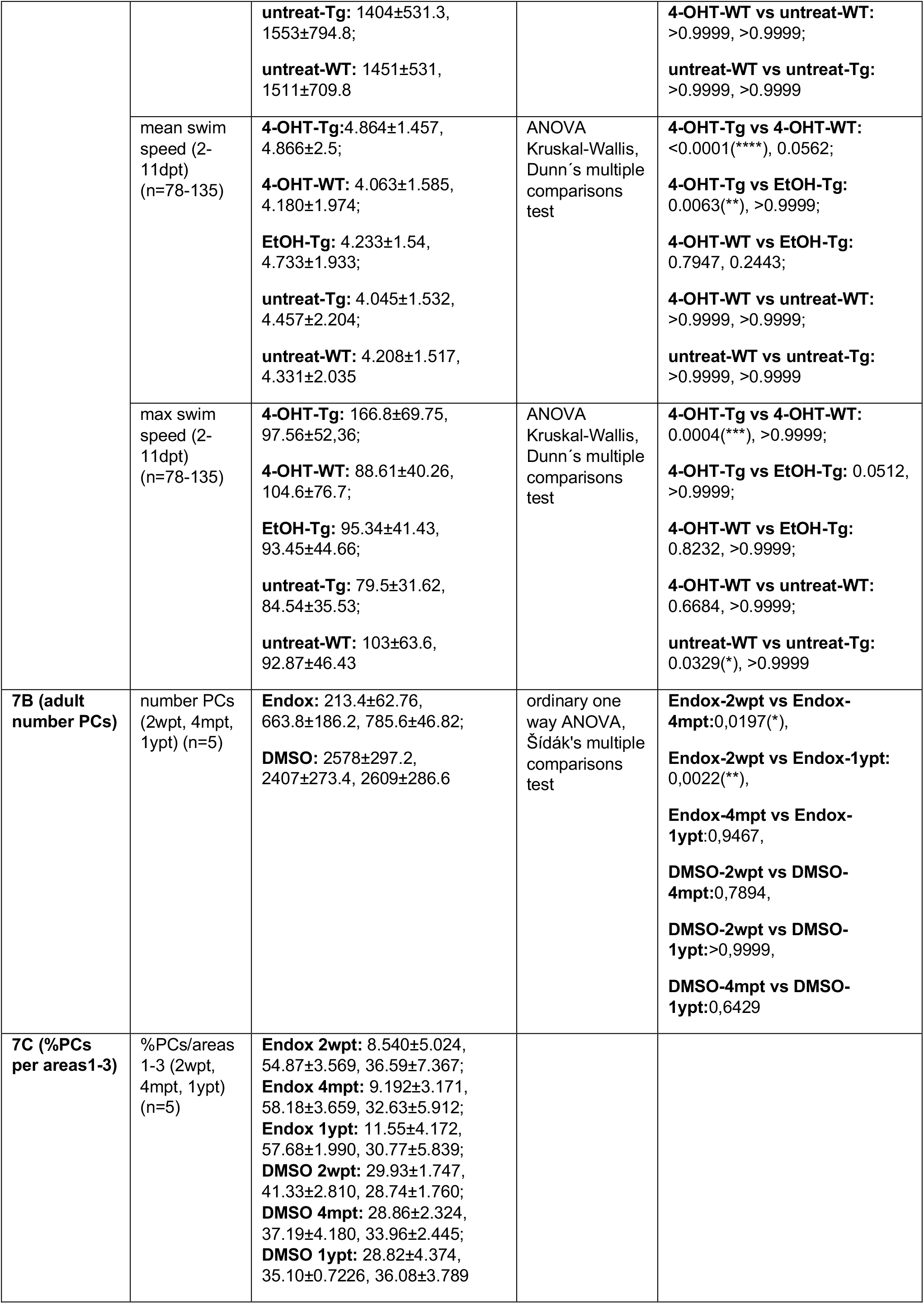
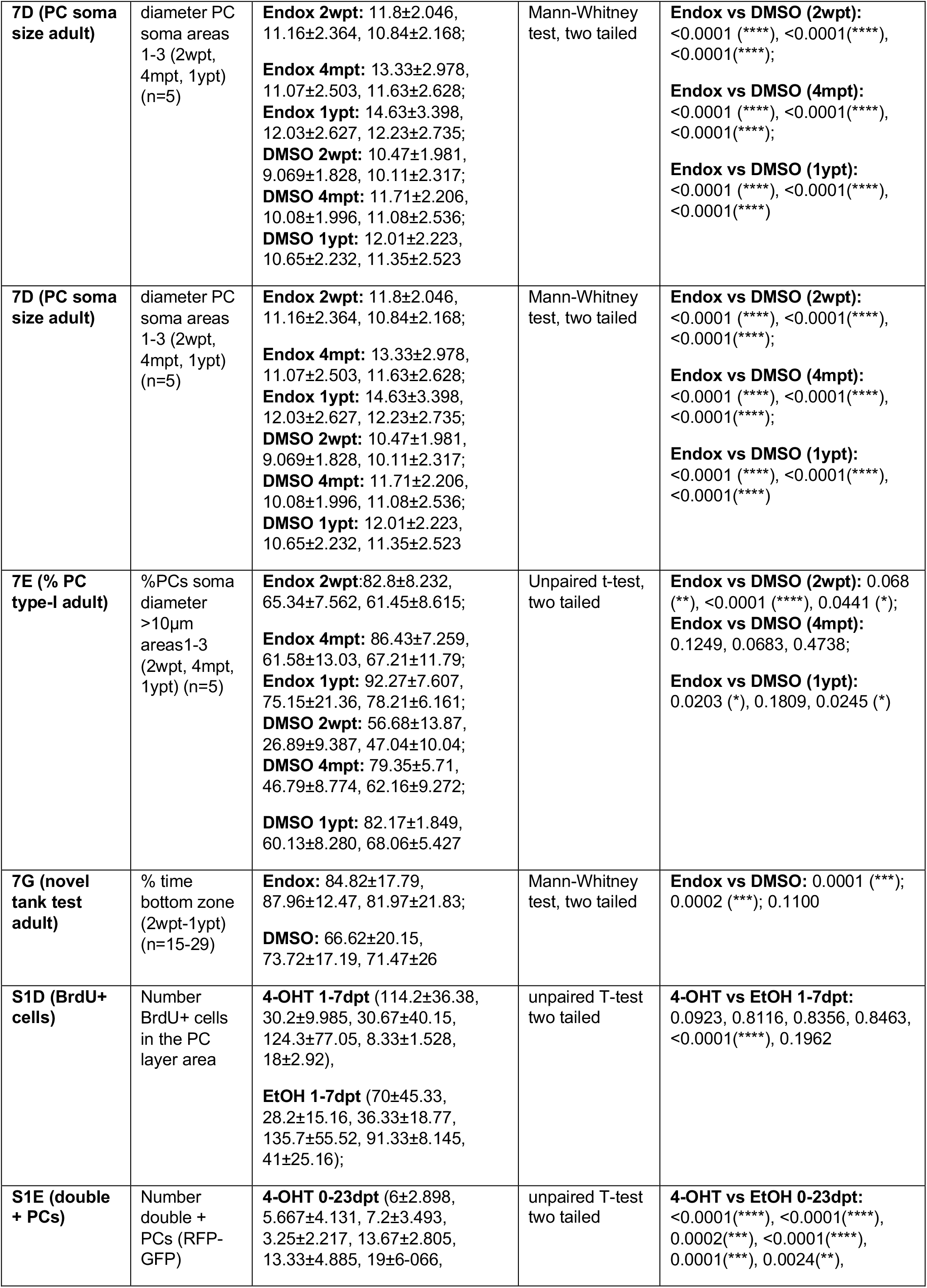
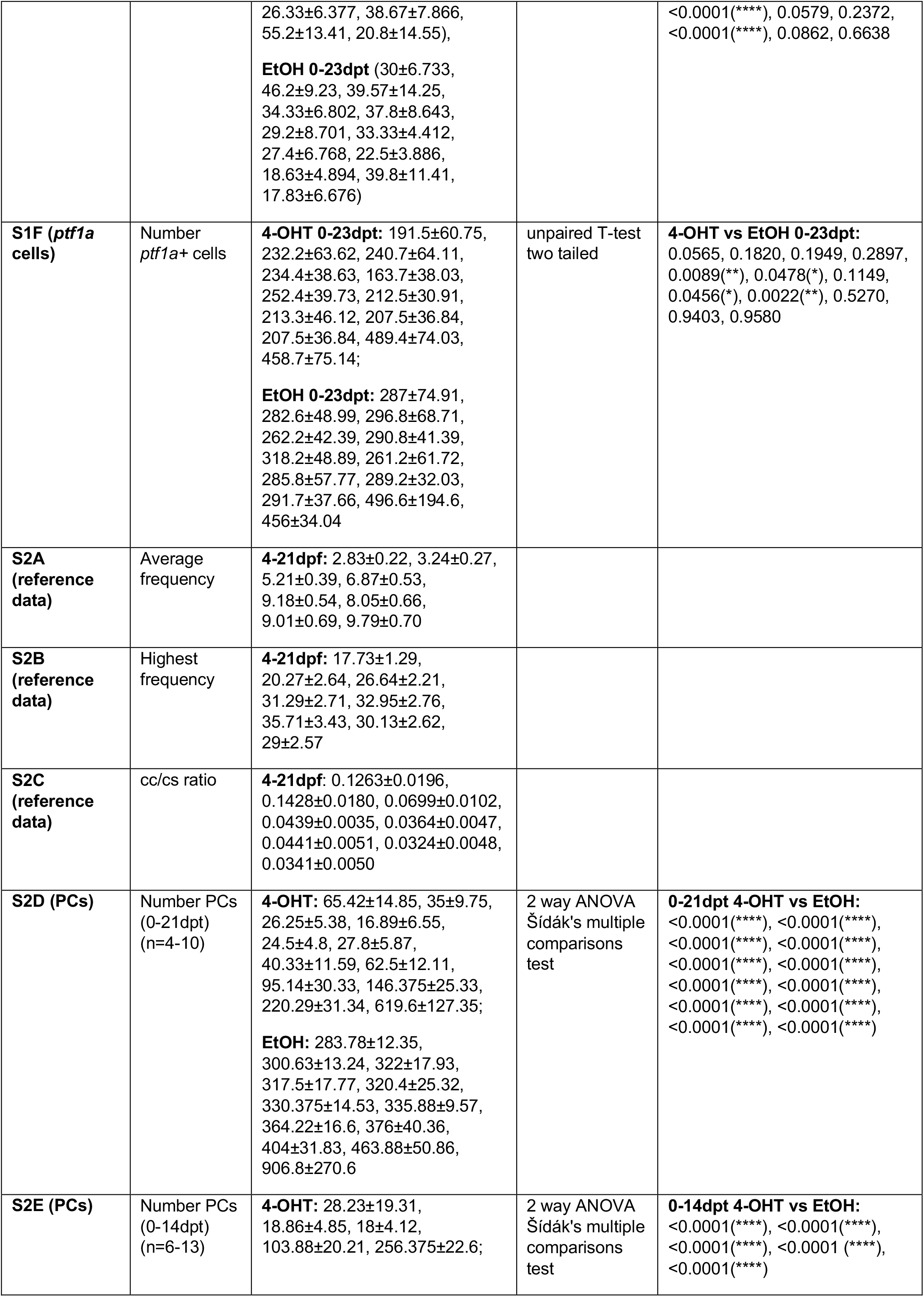
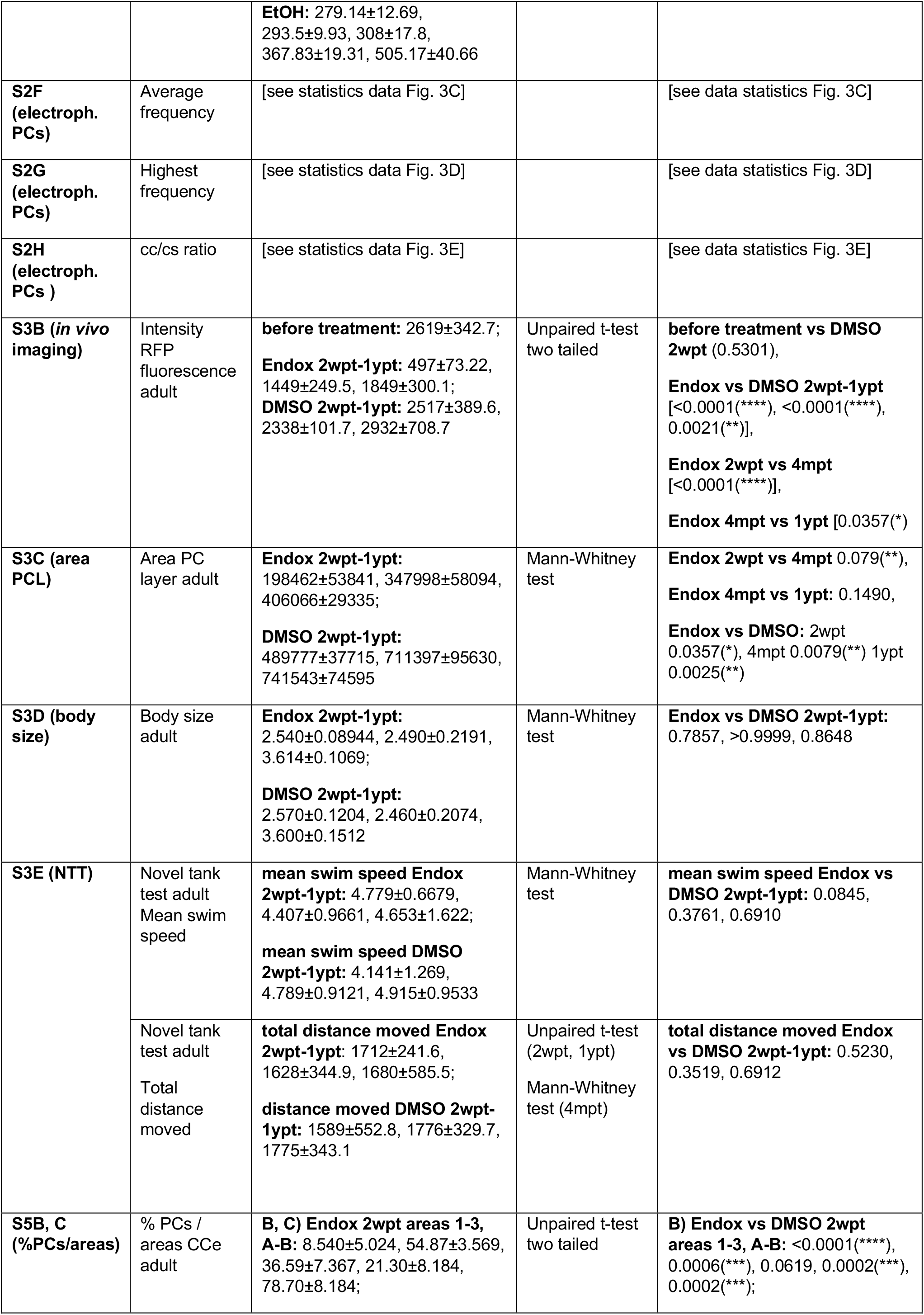
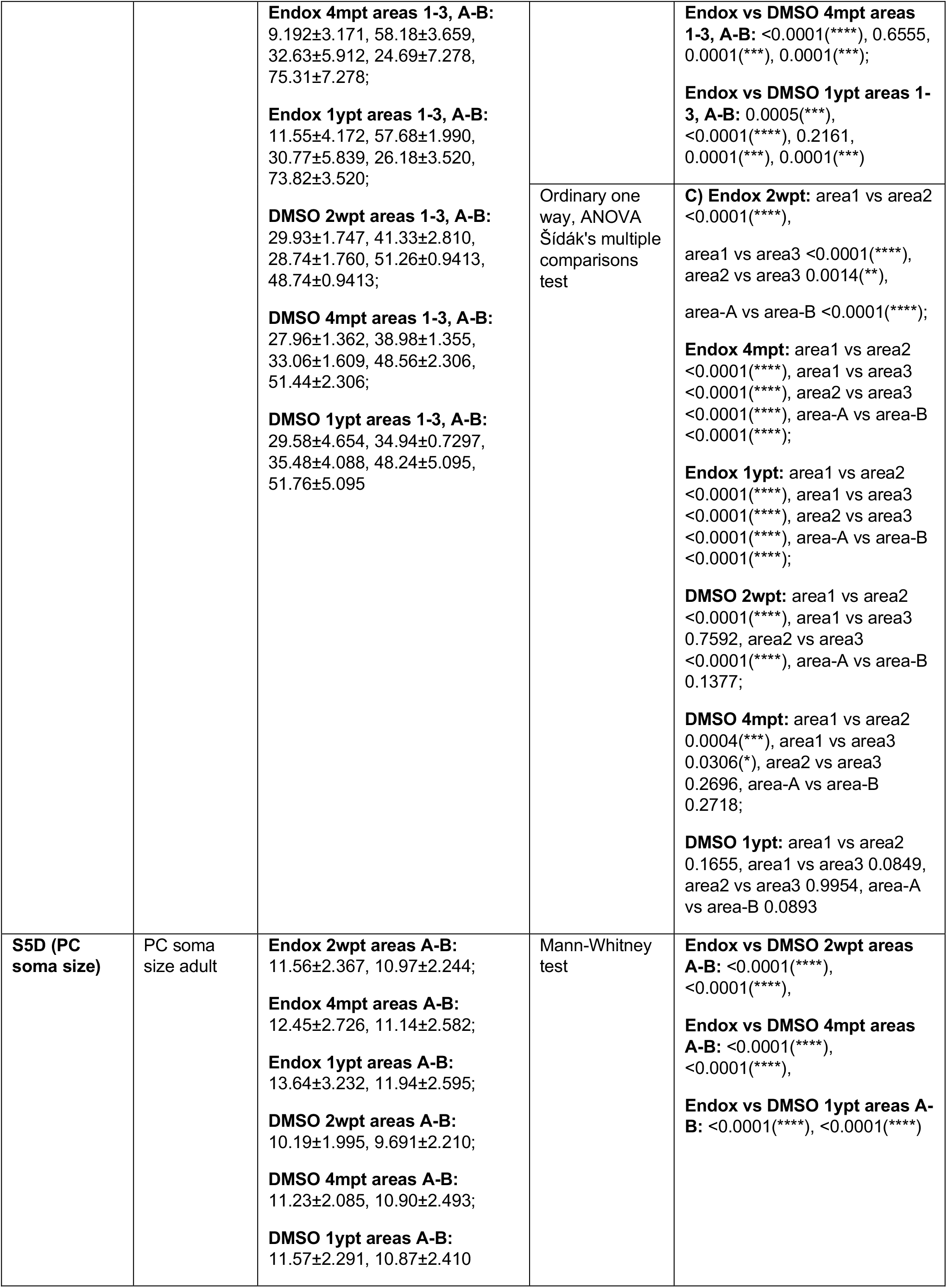
**Table of detailed statistics, related to figures 2-5, 7, and Supplementary Figures 1-3 and 5.** Statistical tests applied, mean standard deviation and n-numbers are indicated, level of significant differences and P values are indicated. Supplementary table 1: statistical data related to the main figures 1-5, 7 and supplementary figures 1-3, 5. Mean, standard deviation, statistic test applied, ‘n’ number, P value and level of significance are indicated. The reference to the different figures and groups of the comparisons performed in each figure are highlighted in bold. Abbreviations: 4-OHT 4-hydroxytamoxifen, BrdU Bromodeoxyuridine, DMSO dimethyl sulfoxide, dpf days post-fertilization, dpt days post-treatment, Endox Endoxifen, EtOH ethanol, *gfap* glial fibrillary acidic protein, mpt months post-treatment, NTT novel tank test, OKR optokinetic response, PC Purkinje cells, PCL Purkinje cell layer, *ptf1a* pancreas associated transcription factor 1a, Tg transgenic, wpt weeks post-treatment, WT wild type, ypt years post-treatment.

**Supplementary Table-2.** Table of key resources and availability.

| REAGENT or RESOURCE |  |  |
| --- | --- | --- |
| Antibodies | Source | Catalog number |
| rabbit anti-tagRFP (1:1.000) | Evrogen, Moscow, Russia | AB233 |
| mouse anti-ZebrinII (1:500) | donated by R. Hawkes, Univ. of Calgary, Canada | N/A |
| chick anti-GFP (1:1.000) | Aves Labs, Davis, CA, USA | GFP-1010 |
| rat anti-BrdU (1:200) | Biozol GmbH, Eching, Germany | Abcam 6326 |
| Alexa goat anti-rabbit IgG (H+L) 568 (1:500) | Invitrogen Inc., Carlsbad, CA, USA | A11011 |
| Alexa goat anti-mouse IgG (H+L) 488 (1:1.000) | Invitrogen Inc., Carlsbad, CA, USA | A11001 |
| donkey anti-chicken IgY (IgG) (H+L) FITC (1:1.000) | Jackson ImmunoResearch, West Grove, PA, USA | 703-545-155 |

| Alexa goat anti-mouse IgG (H+L) 488 (1:1.000) | Invitrogen Inc., Carlsbad, CA, USA | A11006 |
| --- | --- | --- |
| Chemicals, Proteins | Source | Catalog number |
| cis/trans-4-Hydroxytamoxifen (4-OHT) | Sigma-Aldrich, St. Louis, MO, USA | H6278 |
| Endoxifen | Sigma-Aldrich, St. Louis, MO, USA | E8284 |
| BrdU | Sigma-Aldrich, St. Louis, MO, USA | B50021-G |
| d-Tubocurare hydrochlorid-pentahydrat | Sigma-Aldrich, St. Louis, MO, USA | T2379 |
| Tricaine, MS-222 | Sigma-Aldrich, St. Louis, MO, USA | E10521 |
| DAPI | ThermoFisher Scientific, Waltham, MA, USA | 62247 |
| Normal Goat Serum, NGS | Vector Laboratories, Burlingame, CA, USA | S-1000 |
| Albumin Fraction V | Carl Roth GmbH, Karlsruhe, Germany | 80076.2 |
| DMSO | AppliChem GmbH, Darmstadt, Germany | 67-68-5 |

| Experimental Models: Fishlines | Reference | Source |
| --- | --- | --- |
| AB wildtype zebrafish | N/A | fish facility TU Braunschweig |
| *brass* pigmentation mutant zebrafish | N/A | fish facility TU Braunschweig |
| Tg(ca8:-FynTagRFP-T2A-Casp8-ERT2 (PC-ATTAC)<sup>bz11</sup> | (Weber et al., 2016) | fish facility TU Braunschweig |
| Tg(-7.5*ca8*:EGFP)<sup>bz12</sup> | (Kazuhiko Namikawa et al., 2019) | fish facility TU Braunschweig |
| Tg(*olig2*:EGFP)<sup>vu12</sup> | (Shin et al., 2003) | fish facility TU Braunschweig |
| Tg(*ptf1a*:EGFP) | (Godinho et al., 2005) | fish facility TU Braunschweig |
| Tg(*gfap*:EGFP)<sup>mi2001</sup> | (Bernardos and Raymond, 2006) | fish facility TU Braunschweig |
| **Software and Algorithms** | **Source** |  |
| Patchmaster Software | HEKA Elektronik GmbH, Reutlingen, Germany |  |
| IGOR Pro 6.37 | Wavemetrics Inc., Portland, OR, USA |  |
| Leica LAS X | Leica Microsystems GmbH, Wetzlar, Germany |  |
| Prism9 Graph Pad | Graph Pad Software, San Diego, CA, USA |  |
| Excel Microsoft | Microsoft Inc., Redmond, WA, USA |  |
| Free Imaging software FIJI | https://imagej.net |  |
| Ethovision Noldus software | Noldus Inc., Wageningen, The Netherlands |  |
| Corel Draw software | Corel Corporation, Ottawa, Ontario, Canada |  |

